# Six New Species and Two Reinstatements of *Viola* (Violaceae) from China

**DOI:** 10.1101/2025.10.02.679011

**Authors:** Yan-Shuang Huang, Qiang Fan

## Abstract

We describe six new species and reinstate two species of *Viola* (Violaceae) from China, based on material collected in Guangxi, Jiangxi, Hubei, and Chongqing. Field surveys, detailed morphological comparisons, and phylogenetic analyses of ITS and GPI gene sequences placed these taxa within section *Plagiostigma* subsections *Diffusae*. The GPI data provided higher resolution, revealing complex relationships and suggesting ancient hybridization or incomplete lineage sorting in subsect. *Diffusae*. Our findings clarify species boundaries and contribute to understanding the diversity and evolution of *Viola* in China.

## Introduction

The genus *Viola* L. (Violaceae) comprises more than 670 species distributed mainly in temperate and montane regions worldwide (Clausen 1929; Huang et al. 2023b). China is a major center of diversity for this genus, where multiple sections and subsections are represented, notably sect. *Plagiostigma* subsect. *Diffusae* (Marcussen et al. 2022). These species are taxonomically challenging due to high morphological variation and complex evolutionary patterns.

In recent years, botanical surveys across southern China have led to the discovery of many previously unrecognized taxa within *Viola* (e.g., Zhou et al. 2008; Ning et al. 2012; Li et al. 2022). However, the diversity of these subsections remains incompletely documented, particularly in biodiversity-rich provinces such as Guangxi, Jiangxi, Hubei, and Chongqing.

Here, based on comprehensive fieldwork, morphological study, and molecular phylogenetic analyses (ITS and GPI gene sequences), we described six new species and reinstate two species of *Viola* from these regions. Our findings contribute to a better understanding of species boundaries and the evolutionary history of *Viola* in China.

## Materials and methods

We conducted field investigations and observations in Guangxi, Jiangxi, Hubei and Chongqing province during the flowering and fruiting periods. Leaf material of the species were collected and stored in zip-lock plastic bags with silica gel for comparisons and taxonomical treatment, and living specimens were cultivated in the laboratory of Sun Yat-sen University (SYSU). We used a micrometer and a stereomicroscope to observe and measure morphological features to compile detailed descriptions, based on both fresh and dry specimens. Voucher specimens were collected in the Herbarium of Sun Yat-sen University (SYS).

Total DNA was extracted by using the modified CTAB method (Doyle and Doyle 1987). Previously reported primers ITS1 and ITS4 (White et al. 1990) were used to amplify the regions of partial internal transcribed spacer (ITS) 1, 5.8S ribosomal RNA gene and partial internal transcribed spacer 2. PCR amplifications were performed following Fan et al. (2015), sequenced using the Sanger method. The ITS sequences were used to determine the phylogenetic position of the new species within the genus *Viola*. We downloaded the sequences of the species and related ones from NCBI(Table S1)and aligned them using MEGA 6.0 (Tamura et al. 2013) with ClusterW, then manually adjusted the alignment. Phylogenetic inferences were carried out with Maximum Likelihood (ML) using IQ-TREE 2.0.3 (Minh 2020). SYM+I+R2 was selected as the best–fit model according to Bayesian Information Criterion and 2, 000 bootstraps were conducted to evaluate the confidence.

To further resolve the phylogeny of the new species within subsection *Diffusae*, we additionally sequenced and analyzed the GPI gene using high-throughput (next-generation) sequencing. The raw reads were quality-filtered using fastp v0.23.4 (Chen et al., 2018). Filtered sequences were processed with BWA v0.7.18 (Li & Durbin, 2009) for alignment, and SAMtools v1.20 (Danecek et al., 2021) together with BCFtools v1.20 (Danecek et al., 2021) for variant calling and sequence assembly. The GPI sequences were aligned with MAFFT v7.525 (Katoh & Standley, 2013). Phylogenetic analysis was conducted using the same Maximum Likelihood method as for the ITS dataset, with the best-fit model determined as K3Pu+F+G4 according to BIC.

## Results

The Maximum Likelihood tree based on ITS sequences placed the 7 species (5 new and 2 two reinstated species) within *Viola* subsect. *Diffusae*, forming several well-supported clades (bootstrap = 99). Within subsect. *Diffusae*, *Viola lucens* W.Becker, *V. changii* S.Zhou & F.W.Xing*, V. acidophila* Yan.S.Huang et Q.Fan*, V. fluvialis* Yan S.Huang et Q.Fan*, and V. suborbiculata* Yan S.Huang et Q.Fan formed a monophyletic clade (bootstrap = 73). *V. diffusoides* Ching J.Wang *and V. wilsonii* W.Becker grouped together with strong support (bootstrap = 100), and *V. aromatica* Yan S.Huang et Q.Fan *and V. tenuifolia* also clustered together (bootstrap = 99).

To further investigate the phylogenetic relationships within subsect. *Diffusae*, we conducted Maximum Likelihood analysis based on GPI gene sequences. Following Marcussen et al. (2022), the GPI data were analyzed separately for two subgenomes to elucidate evolutionary histories, the GPI tree revealed a clearer phylogeny compared to the ITS tree. Notably, *V. orientosinensis* Yan S. Huang, Y.Xiong & Q.Fan clustered closely with *V. qingruii* Yan S.Huang & Q.Fan, and *V. amamiana* Hatus., rather than being sister to *V. nanlingensis* J.S.Zhou & F.W.Xing. *V. nanlingensis* was found to be non-monophyletic; however, a clear phylogenetic affinity was observed between *V. suborbiculata* and this species. *V. diffusoides, V. wilsonii, V. aromatica, and V. tenuifolia* showed close relationships with *V. diffusa* and systematic position of *V. fluvialis* showed considerable conflict between the two subgenomes. *V. acidophila* clustered together with specimens of *V. changii* from different localities, with *V. changii* forming a monophyletic group. These two species were recovered as well-supported sister clades (bootstraps = 100; 100)

### Taxonomic treatment

***Viola acidophila*** Yan S. Huang & Q. Fan, sp. nov. (Fig. 3, 4)

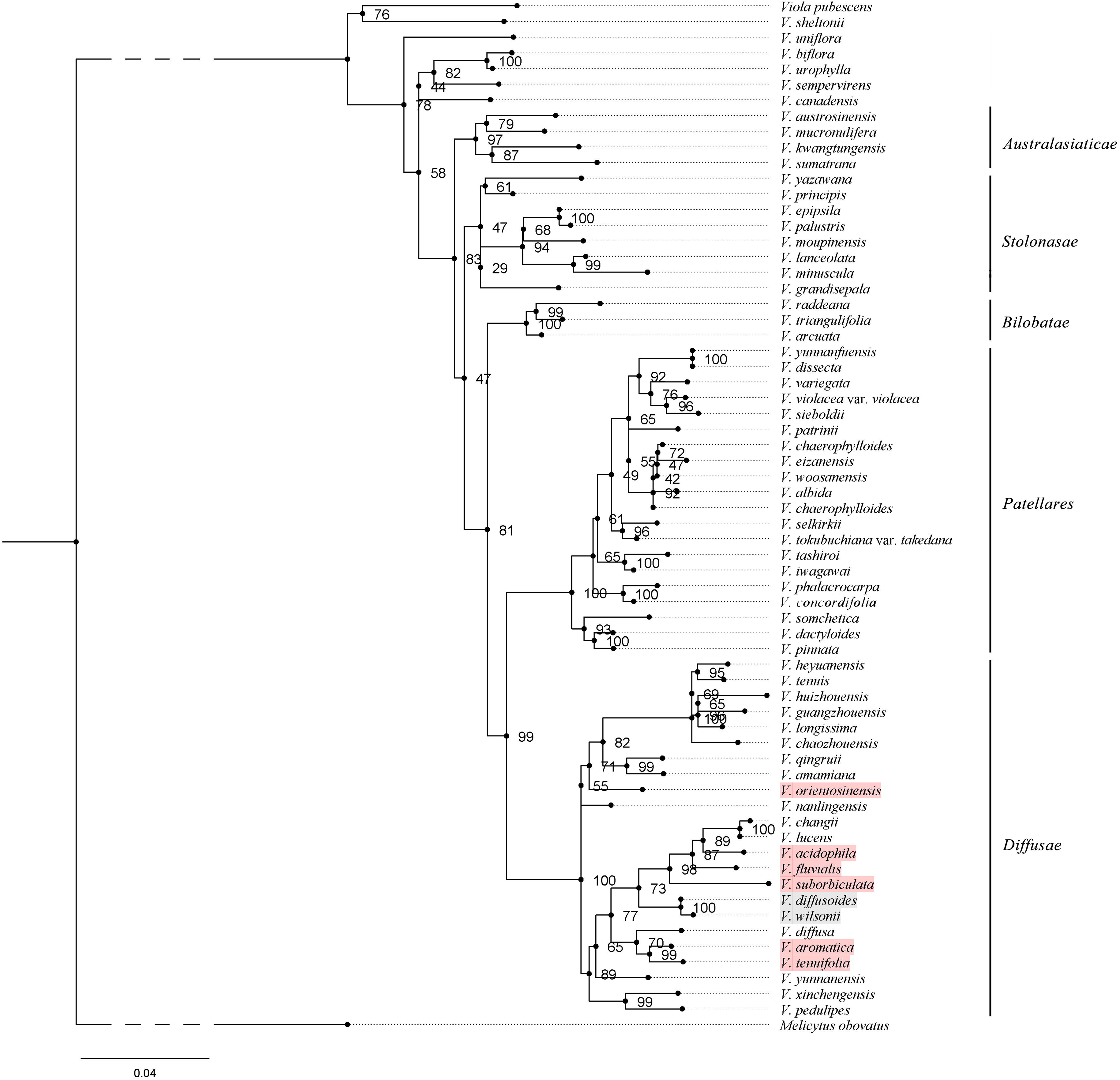

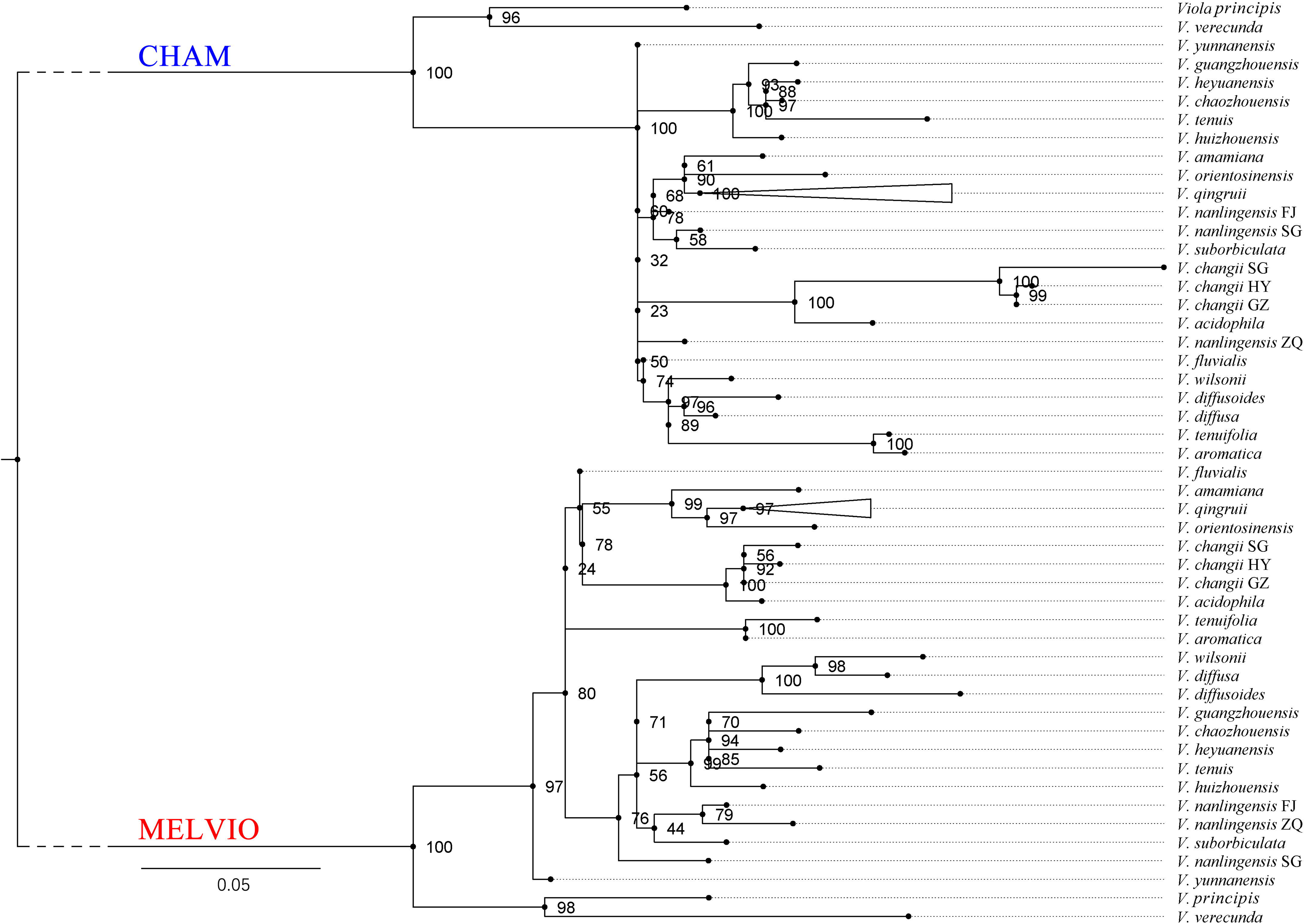

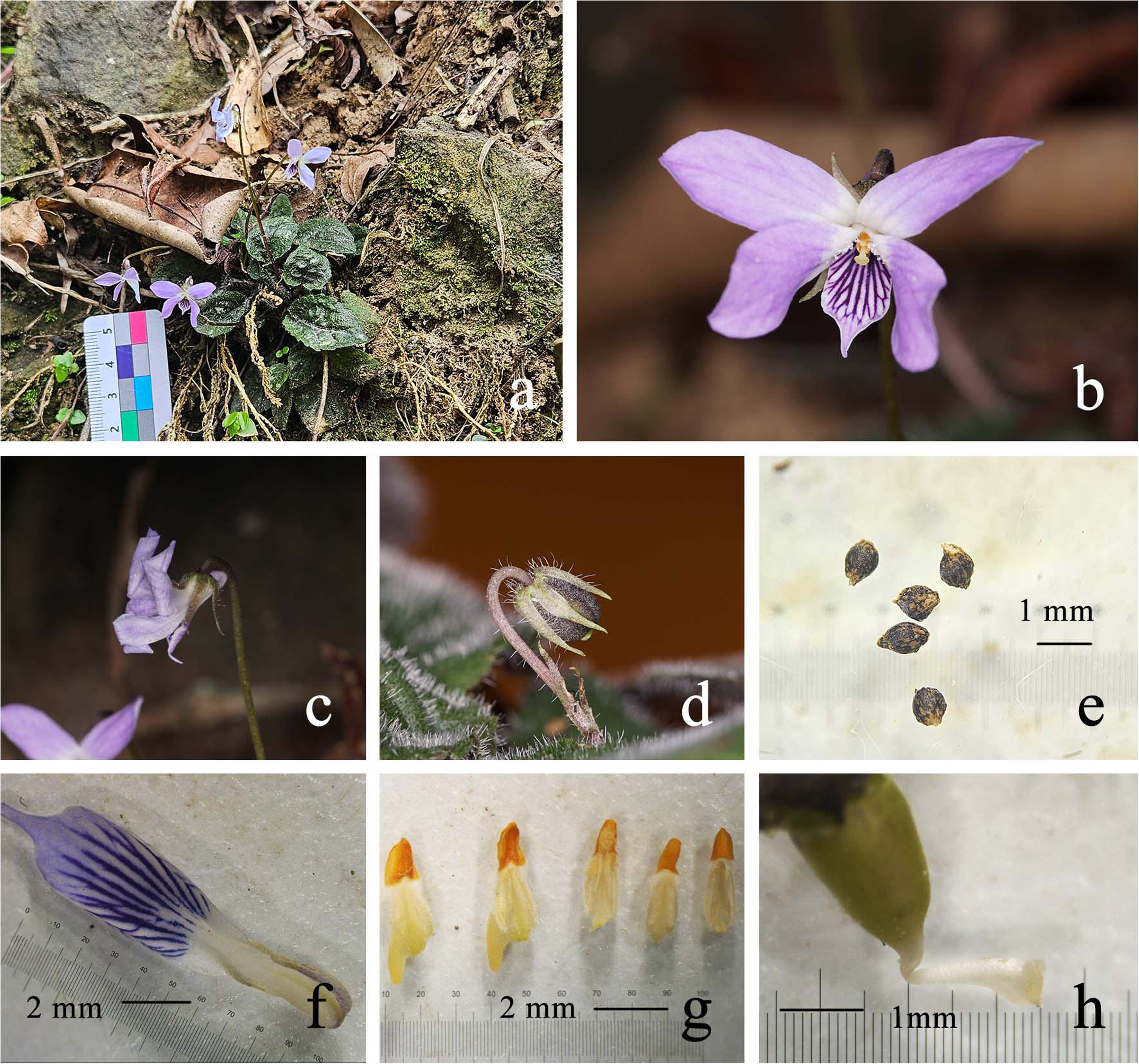

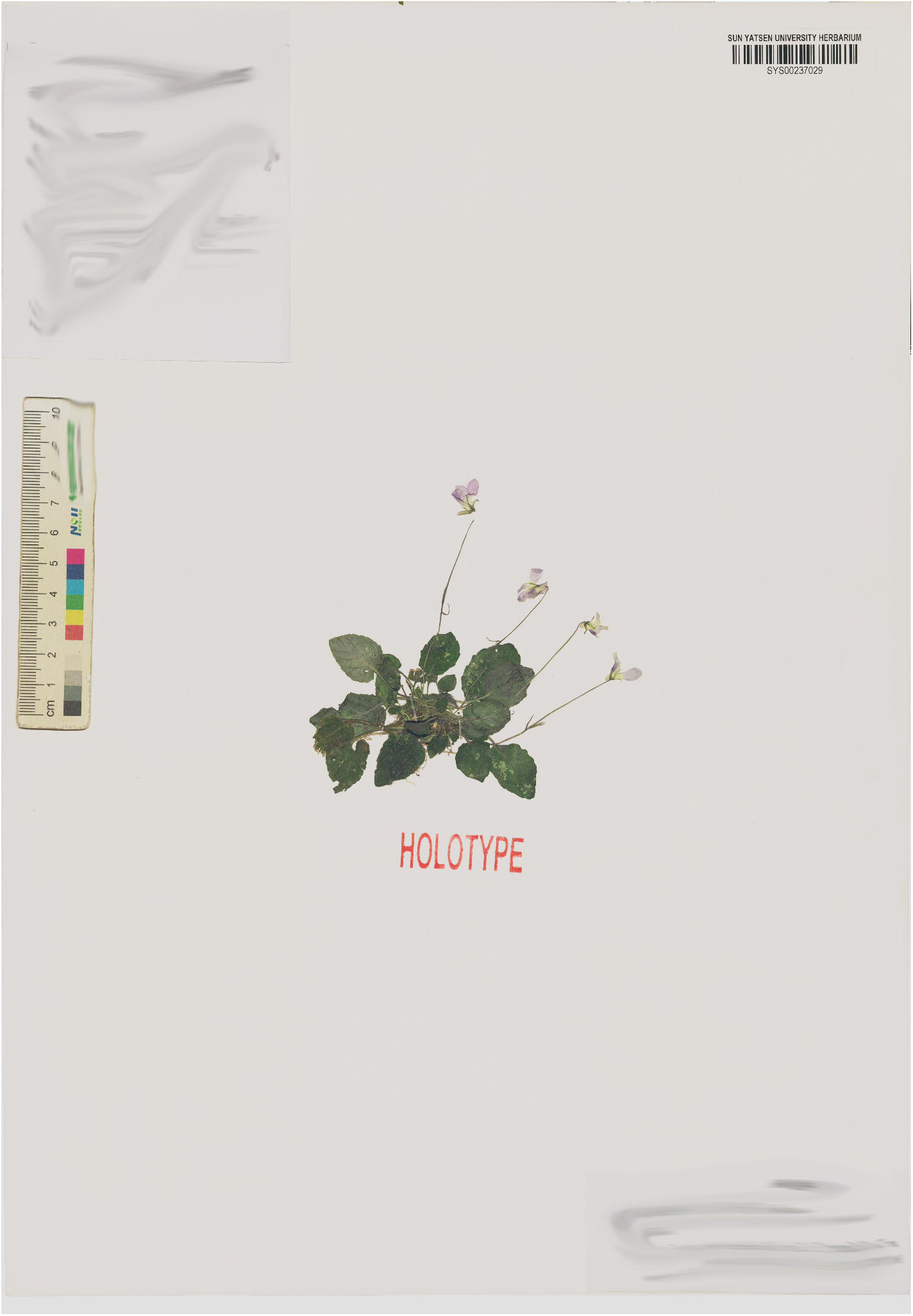

#### Diagnosis

A species morphologically close to *Viola changii*, but differing in having oblong leaves, densely puberulent, with an acuminate apex and dull surface (vs. ovate leaves, glabrous or with rough hairs, with apex obtuse and glossy surface); seeds densely tuberculate (vs. seeds relatively glabrous and smooth); stolons absent and reproduction by tillering (vs. presence of obvious lateral stolons).

**Type:** China. Guangxi: Guilin City, Lipu, growing in cool valleys and along forest edges on acidic red and yellow soils, 24°42′36.98″ N, 110°11′18.3″ E, 382 m a.s.l., 26 Mar. 2025, *Yan. S. Huang HYS25032601*(holotype: SYS00237029).

***Additional specimens examined:*** China. Guangxi, Yangshuo, 21 Sep. 2018, *450321180921042LY* (LBG00414024); Guangxi, Yongfu, 23 Mar. 2009, *Yingfeng Huang* et *Xingmo Luo 20126* (GXMG0010061); Guangxi, Lipu, 27 Oct. 2018, 450331180827027LY (IBK00415411); Guangxi, Pingle, 15 June 2018, 450330180515029LY (IBK00412188); Guangxi, Ziyuan, 4 April 2025, *Hong Zhang* 01116 (PE01334674); Hunan, Yongzhou, 16 April 2020, *Xiong Li*, *Ang Liu* et *Youke Gong* LK0342 (CSFI073335).

#### Etymology

The specific epithet refers to the species’ distribution in the red and yellow soil areas surrounding the karst region.

#### Description

Perennial, tillering herb with basal leaves rosulate, ca. 7 cm tall at flowering and lying on the ground when not flowering. Rhizomes erect or ascending, slender, and very short. Stolons absent; Lateral stems ascending, each terminating in a rosette of leaves at the apex. Stipules 3–5 mm long, adnate to petioles at base, lanceolate, apex acute, pubescent along the margin. Petioles ca. 1.5 cm, densely pubescent along the margin. Leaf blades oblong, 1.6 – 2.0 × 2.4 – 3.2 cm, puberulent; margin serrate; base cordate while living; apex acuminate. Chasmogamous flowers ca 1.5 cm in diam.; peduncles slender, 5–6 cm long, pubescent at about 1/3 base, rising well above the leaves, with two opposite bracteoles above middle; bracteoles linear-lanceolate, 6 mm long, apex cute, pubescent. Sepals green, with densely purple spots, pubescent, lanceolate, 1.1 – 1.3 × 5.5 – 5.0 mm, apex acute, base truncate, appendages absent. Petals purple, the anterior one with apparent violet lines; posterior petals narrowly obovate, 4.0 × 8.5 mm, glabrous, with entire margin and obtuse apex; lateral petals with straight to slightly clavate hairs at the base, oblong, 3.5 × 11 mm, with entire margin and obtuse or erose apex; anterior petal spatulate, apex acute, with a short saccate spur at base, with interior side of base puberulent, including spur 1.3 cm long. Stamens 5, unequal, puberulent; anther thecae 1.5 – 1.8 mm long, with terminal appendages 0.6 – 0.8 mm long; posterior appendages (nectar spurs) of the two anterior stamens ca 1.5 mm long, triangular. Ovary ovoid to ellipsoid, ca. 1.6 mm, glabrous; style clavate, ca 1.6 mm long, conspicuously geniculate at base; stigma glabrous, with thickened lateral margins and a membranous apex. Cleistogamous flowers in June ca. 4 mm long; peduncles 1.2 – 2.5 cm long; bracteoles linear-lanceolate, 6 mm long, acuminate at apex. Capsules puberulent, dark purple, ovoid to oblongoid, 4 – 5 mm long. Seeds nearly ovate, 1 mm long, dark brown, surface densely covered with tubercles; elaiosomes absent.

#### Phenology

Chasmogamous flowers from March to April, cleistogamous flowers from May to September, and fruits from April to November.

#### Distribution and habitat

This species occurs in the deep valleys surrounding the karst landscapes of Guilin, growing along forest margins on red or yellow soils.

#### Conservation status

Each known population consists of fewer than 50 individuals. Due to its restricted distribution, small population size, and observed decline in habitat quality, the species is assessed as Endangered (EN) under the IUCN Red List criteria(B1; B2)

***Viola aromatica*** Yan S. Huang et Q. Fan, sp. nov. (Fig. 5, 6)

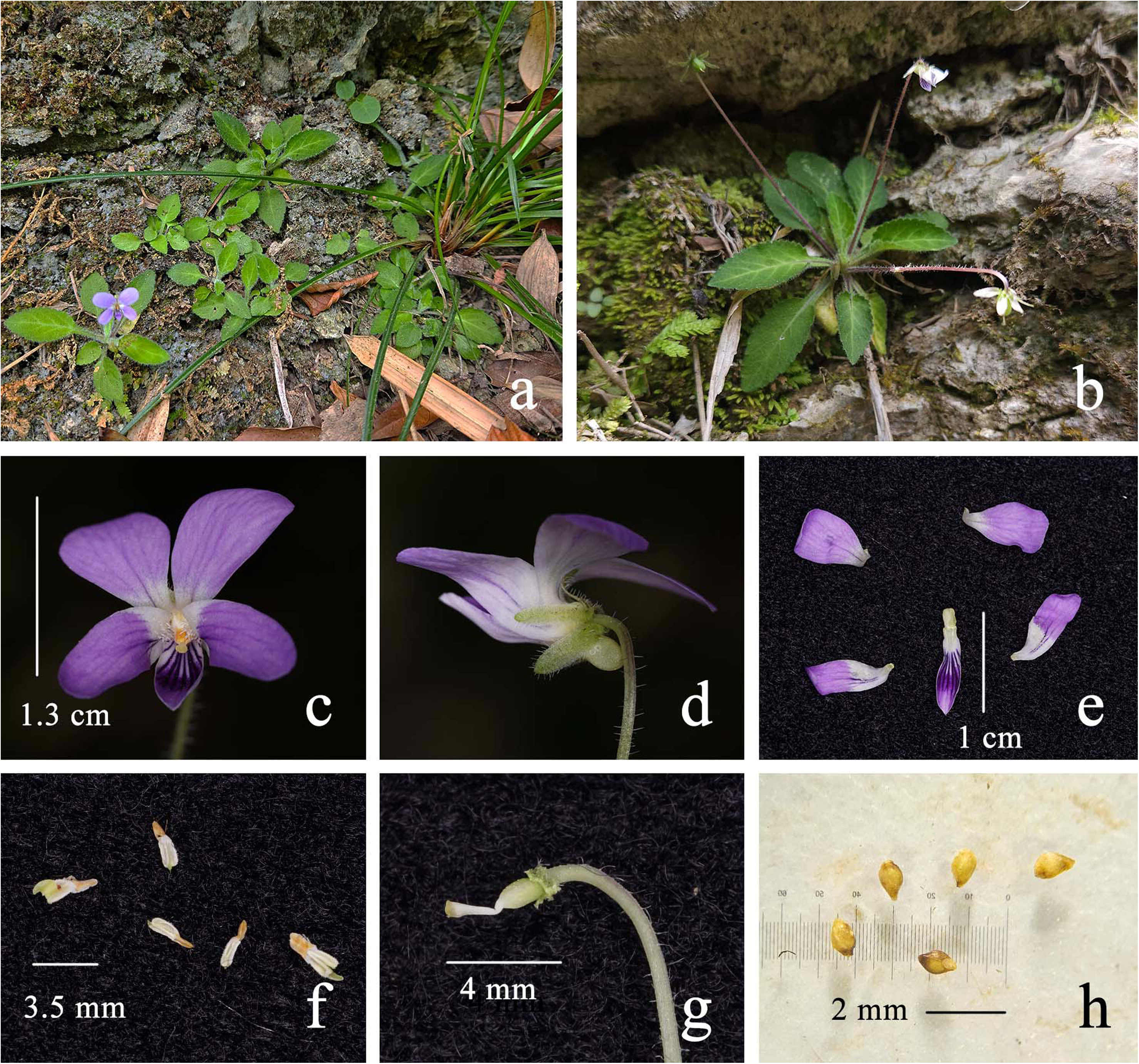

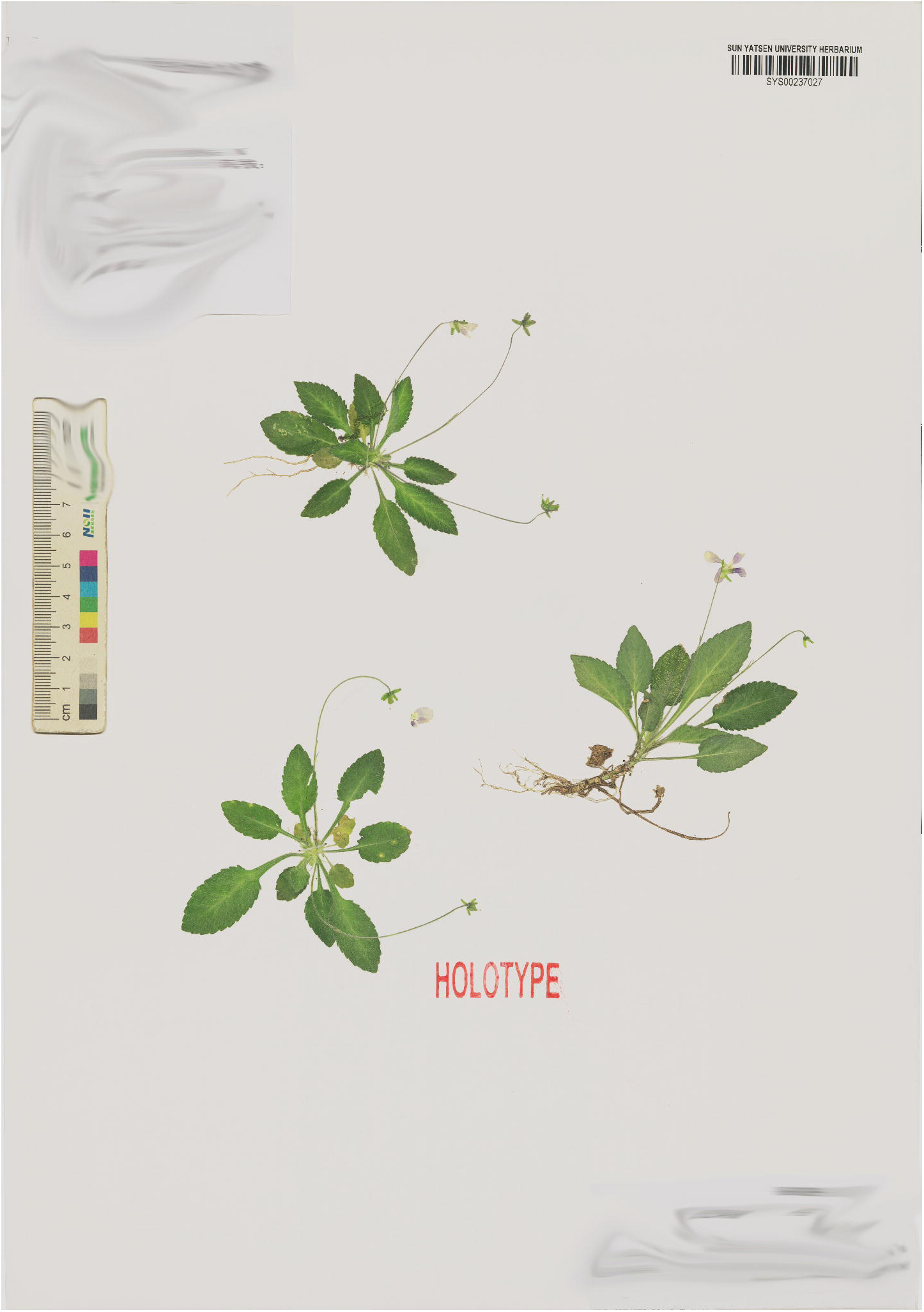

#### Diagnosis

This species is morphologically similar to *Viola diffusa* Gings, but differs in having leaves densely puberulent (vs. leaves sparsely scabrous-pubescent), larger flowers with deeper coloration (vs. smaller, pale pink flowers), lateral petals without distinct yellow spots at the base (vs. lateral petals with conspicuous yellow to green spots), and flowers strongly fragrant (vs. flowers without obvious scent).

***Type*:** China. Chongqing: Beibei District, in rocky crevices near the valley, 30°0′50.22″ N, 106°37′15.64″ E, 500 m a.s.l., 16th. April, 2025, *Yan. S. Huang HYS25041601* (holotype: SYS00237027).

#### Etymology

The specific epithet refers to its strong fragrance.

#### Description

Annual or perennial herbs; basal leaves rosulate; plants ca. 7 cm tall at flowering, otherwise 3 – 4 cm. Rhizome absent or ascending, slender, ca. 1.5 cm. Lateral stems or stolons absent. Stipules 4 – 6 mm, adnate to petioles at about 1/3 base, lanceolate, apex acute, margin fimbriate, puberulent. Petioles 1.2 – 1.8 cm long, the leaf base decurrent along the petiole forming narrow wings, puberulent. Leaf blades ovate to oblong, 1.1 – 1.4 × 1.8 – 3.2 cm, puberulent, apex acuminate to obtuse, margin serrate, base truncate or decurrent. Chasmogamous flowers 2.1 cm in diam.; Peduncles 4.5 – 6.5 cm long, puberulent; with a pair of bracteoles at the middle; bracteoles linear-lanceolate, ca. 5 mm long, apex acute, puberulent. Sepals green, pubescent, lanceolate, 1 – 1.5 × 3.8 mm, apex obtuse, base with shortly round appendages. Petals purple, the anterior one with apparent violet lines; posterior petals obovate, 4.5 × 8.0 mm, glabrous, with entire margin and erose apex; lateral petals with straight to slightly clavate hairs at the base, oblong, 3.5 × 10 mm, with entire margin and erose apex; anterior petal narrowly oblong, with entire margin and acuminate apex, and a short saccate spur at base, interior side of base puberulent, including spur 1.0 cm long. Stamens 5, unequal, puberulent; anther thecae 1.5 mm long, with triangular terminal appendages 0.8 – 1.5 mm long; posterior appendages (nectar spurs) of the two anterior stamens 1.3 mm long, triangular. Ovary ovoid, 1.7 mm, puberulent; style clavate, 1.8 mm long, conspicuously geniculate at base; stigma glabrous, with thickened lateral margins and a membranous apex. Capsules puberulent, green ovoid to oblongoid, 3 – 4 mm long. Seeds yellow, ovoid, ca. 0.8 mm long, glabrous; elaisomes inconspicuous.

#### Phenology

Chasmogamous flowers in April, cleistogamous flowers started from May, and fruits from April to September.

#### Distribution and habitat

*Viola aromatica* is currently only known from Jindao Cyon, grows along the valleys.

#### Conservation status

Due to the lack of further surveys in surrounding areas, the conservation status of this species is assessed as Data Deficient (DD) according to the IUCN criteria.

***Viola diffusoides*** Ching J. Wang, *Bull. Bot. Res. Harbin* 8(2): 19. 1988. stat. rev. (Fig. 7)

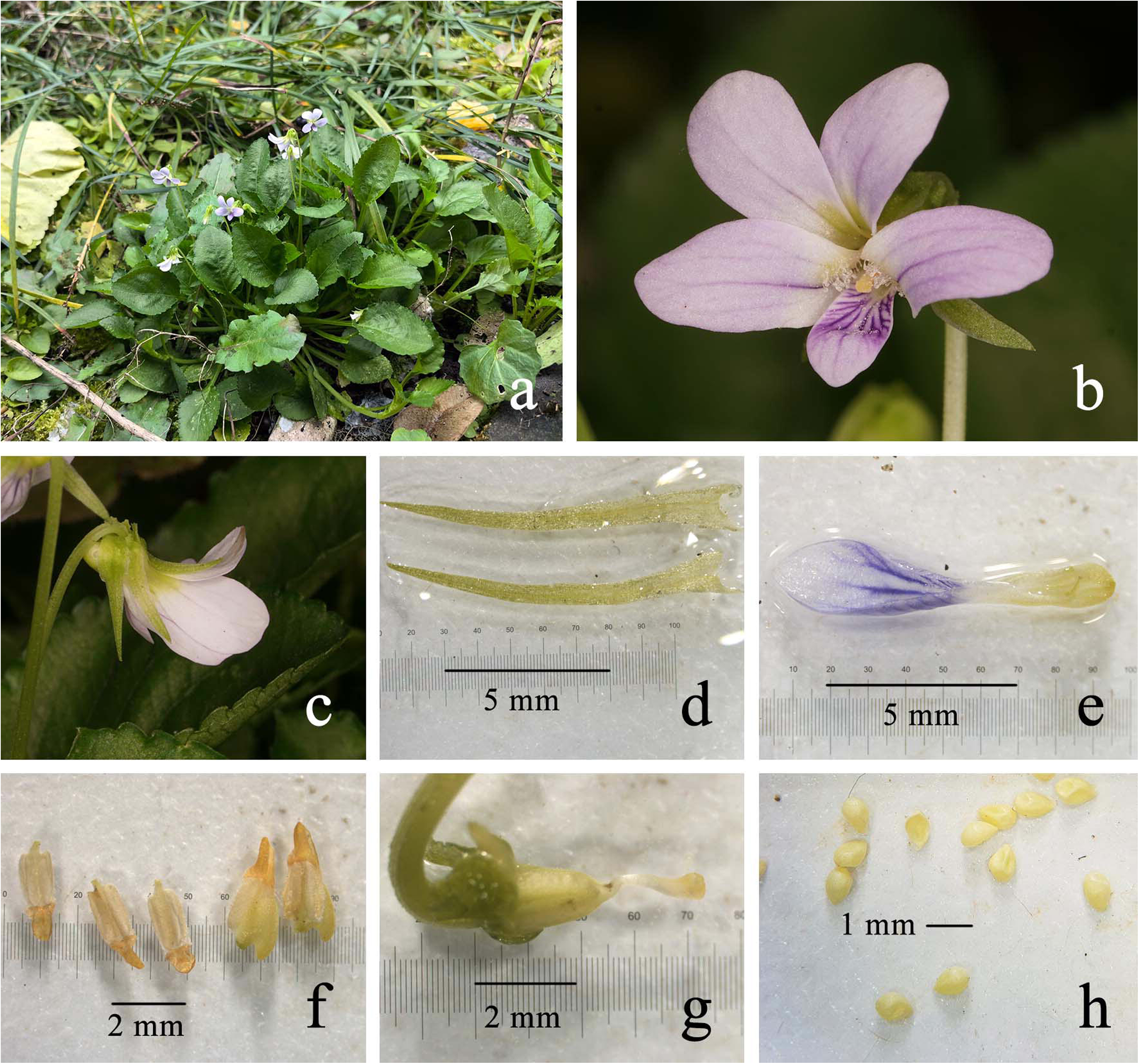

#### Diagnosis

This species is morphologically similar to *Viola diffusa* Gings, but differs by being completely glabrous, having a well-developed rhizome.

***Type:*** China. Sichuan: Hehuan, between Jintan and Quhekou, on mountain slopes, 19 March 1941, *Y.C. Yang 4231* (holotype: PET00025459).

***Additional specimens examined:*** China. Chongqing, Beibei Dongyang town, 15 Feb. 2024, *Yan. S. Huang HYS24021501* (SYS00237031). Chongqing, Beibei, 1 July 1956, *Chuanqiandui* 502(PE 02083260; Chongqing, Beibei, 14 Nov. 2018, *Yi Yang IMPCYY019B09* (KUN 1550624); Sichuan, Tianquan, 14 Sep. 1963, Kejian Guan et Wencai WANG 3449 (PE00067136); Sichuan, Tianquan, 14 Sep. 1963, Kejian Guan et Wencai WANG 3449 (PE00067135); Guizhou, Zhengan, 17 Mar. 2013, *Hong Liu 520324160317009LY* (GZTM0043677).

#### Description

Perennial herbs, aerial stem absent. Rhizomes erect, ca. 8 mm long, 1 – 8 mm in diam., rigid, green. Stolon slender and creeping. Basal leaves rosulate, or alternate on the stolons; leaf blades ovate to oblong, 0.8 – 1.5 × 1.5 – 2.5 cm, apex obtuse or slightly acuminate, base round or truncate, decurrent to the petioles, margins finely crenate, glabrous; petioles 1.0 – 3.5 cm long, with narrow wings, glabrous; stipules 1 – 1.5 cm long, 1.5 mm wide, only adnate to petioles at base, linear-lanceolate, apex acuminate, margin sparsely fimbriate, glabrous. Chasmogamous flowers small, pale violet, with slender pedicels; Peduncles arise from the basal leaves or the leaf axils of creeping stems, 3 – 8 cm long, glabrous, with a pair of bracteoles above the middle; bracteoles linear-lanceolate, 6 mm – 12 long, margins entire; sepals lanceolate, 6 – 7.5 mm long, apex gradually acuminate, with white membranous margins, basal appendages short, about 1.5 mm long, claw-like; the anterior one with apparent violet lines; posterior petals oblongly ovate, 5 – 9.5 mm long, glabrous, with entire margin and round apex, base gradually narrow; lateral petals with straight to slightly clavate hairs at the base, oblong, 5 – 9.5 mm long, apex round; anterior petal narrowly oblong, with entire margin and acuminate apex, and a short saccate spur at base, interior side of base puberulent, including spur 7.5 – 9 mm long. Stamens 5, unequal, puberulent; anther thecae 1.6 mm long, with triangular terminal appendages 0.8 – 1.0 mm long; posterior appendages (nectar spurs) of the two anterior stamens 1.2 – 1.4 mm long, triangular. Ovary ovoid, 1.8 mm, glabrous; style clavate, 1.8 mm long, conspicuously geniculate at base; stigma glabrous, with thickened lateral margins and a membranous apex. Capsules glabrous, green, oblongoid, 6 – 7 mm long. Seeds yellow, ovoid, ca. 0.8 mm long, glabrous; elaisomes inconspicuous.

#### Distribution and habitat

According to the original description, this species is distributed in Chongqing, Sichuan, and Yunnan, China. The type locality is on mountain slopes.

#### Conservation status

Due to its wide distribution, this species is assessed as Least Concern (LC) according to the IUCN criteria.

#### Notes

This species was treated as a synonym of *Viola diffusa* by Chen in *Flora of China* Vol. 13 (2007). Based on morphological and molecular evidence, we here reinstate it as a distinct species.

***Viola fluvialis*** Yan S. Huang et Q. Fan, sp. nov. (Fig. 8, 9)

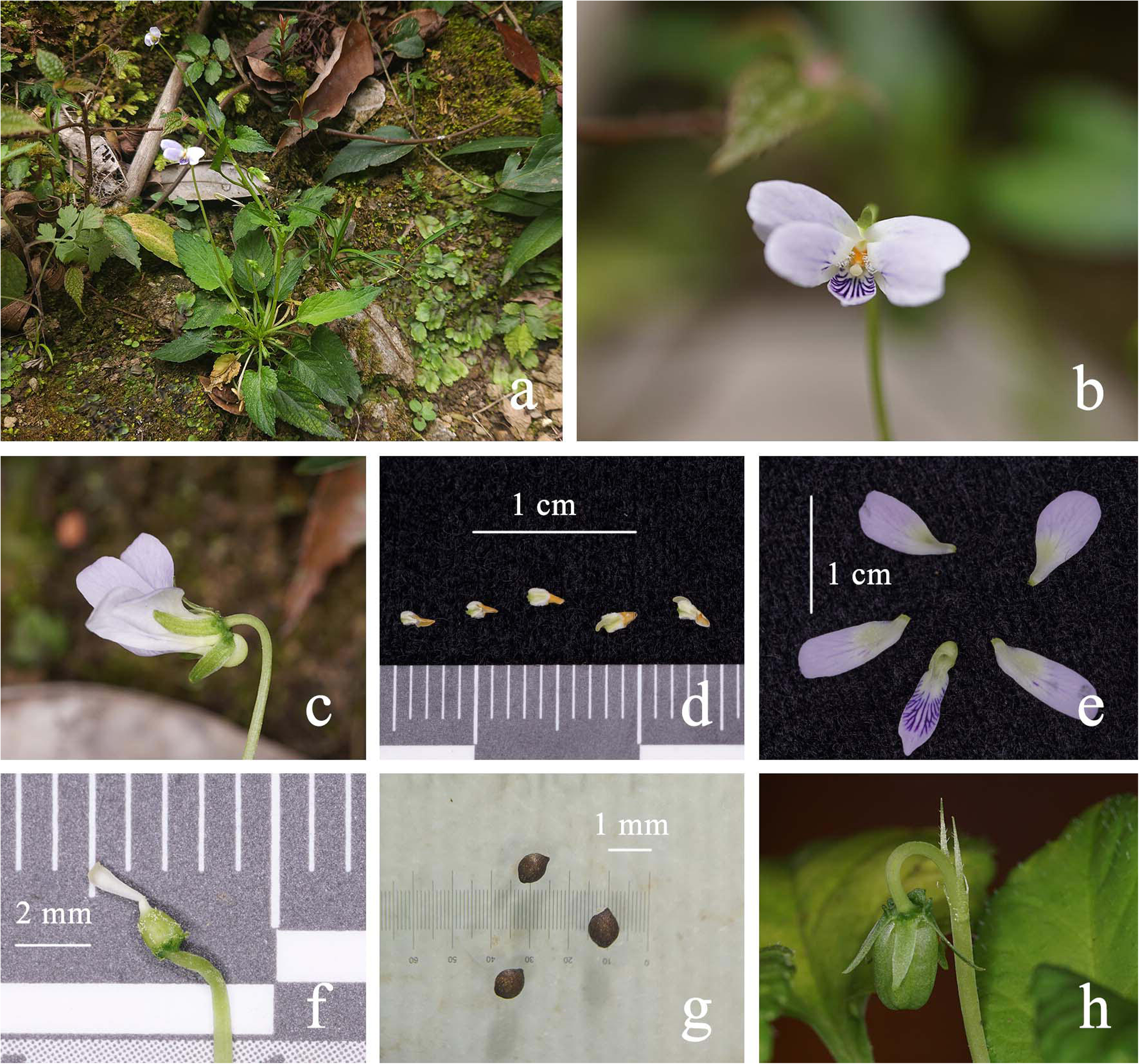

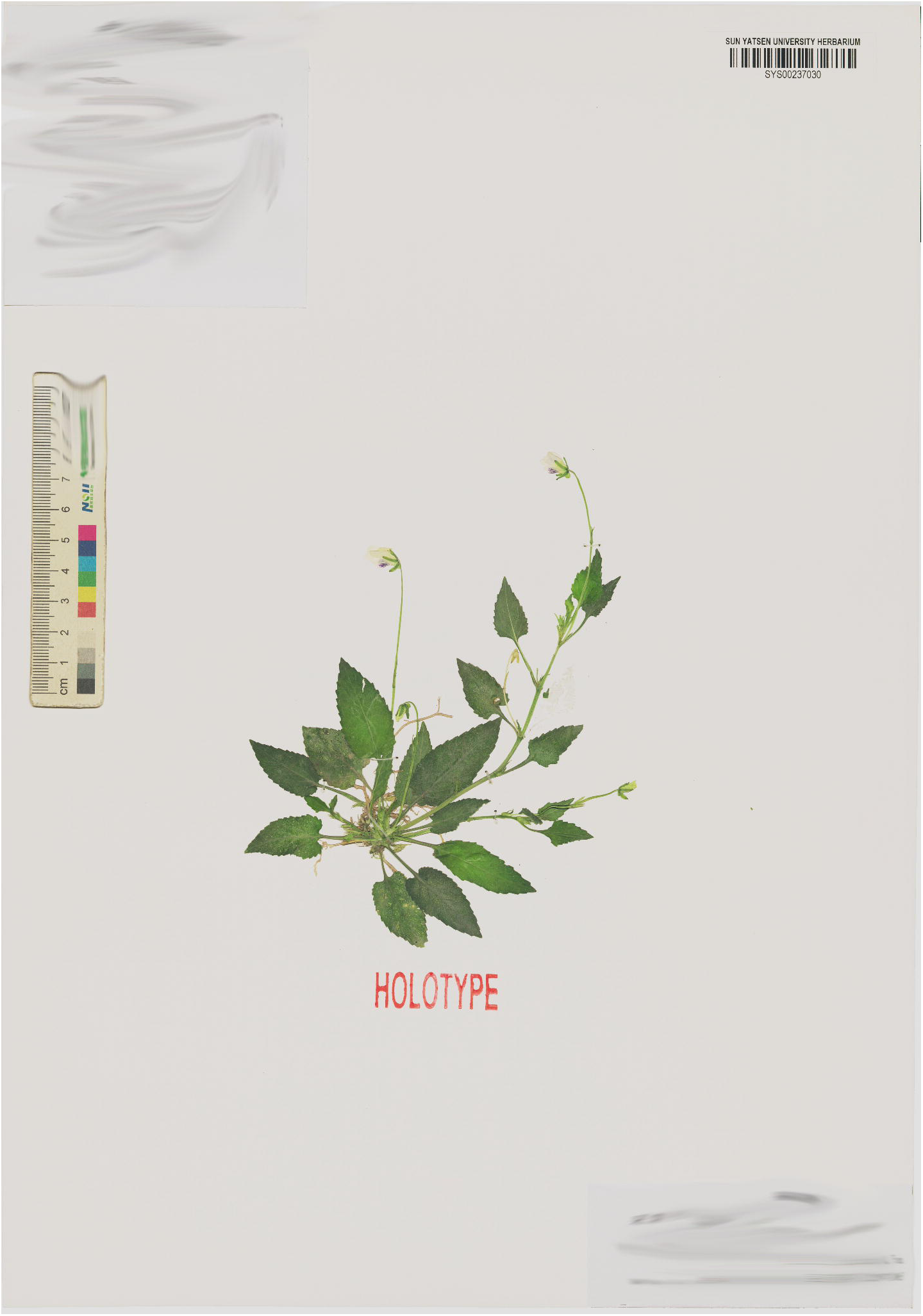

#### Diagnosis

This species is morphologically close to *Viola yunnanensis* W.Becker & H. Boissieu, but differs in having sparsely pubescent leaves and veins that are not purple (vs. densely hirsute leaves with veins usually purple), stigma not shortly beaked in front (vs. stigma shortly beaked in front), and seeds dark brown (vs. yellow seeds)

***Type*:** China. Guizhou: Tongren City, Songtao County, in rocky crevices near the valley, 27°52′58.31″ N, 109°15′3.59″ E, 450 m a.s.l., 2 April 2025, *Yan. S. Huang HYS25040201* (holotype: SYS00237030).

***Additional specimens examined:*** China, Guangxi, 11 July 2012, *Richeng Peng, Shuisong Mo* et Jing Liu ML 1466 (IBK00309696).

#### Etymology

The specific epithet refers to its habitat by streams.

#### Description

Annual herbs, basal leaves tufted, 7 – 10 cm tall when flowering. Rhizome short and creeping, ca. 1 cm. Lateral stems ascending, ca. 12 cm tall, leaves produced at the nodes, with flower buds forming in the leaf axils. Stipules 4 – 7 mm, adnate to petioles at about 1/3 to 1/2 base, triangular to lanceolate, apex acute, margin fimbriate, glabrous. Petioles 1.1 – 1.9 cm long, the leaf base decurrent along the petiole forming narrow wings, glabrous. Leaf blades triangular to oblong, 1.1 – 1.7 × 2.1 – 3.5 cm, apex acuminate, margin serrate, base truncate or cordate, sparsely pubescent along green veins. Chasmogamous flowers ca. 1.2 cm in diam.; Peduncles 4.8 – 8.5 cm long, glabrous; with a pair of bracteoles at the middle; bracteoles linear-lanceolate, 4.5 mm long, apex acute, margin sparsely serrate, glabrous. Sepals green, glabrous, lanceolate, 1.5 × 3.5 – 4.5 mm, apex acuminate, base with short, triangular to claw-like appendages. Petals purple, the anterior one with apparent violet lines; posterior petals obovate, 3.5 × 7.0 mm, glabrous, with entire margin and erose apex; lateral petals with straight to slightly clavate hairs at the base, oblong, 2.5 × 8.0 mm, with entire margin and erose apex; anterior petal narrowly oblong, with entire margin and acuminate apex, and a short saccate spur at base, interior side of base puberulent, including spur 7.5 mm long. Stamens 5, unequal, puberulent; anther thecae 1.0 – 1.5 mm long, with triangular terminal appendages 0.8 – 1.2 mm long; posterior appendages (nectar spurs) of the two anterior stamens 1.5 mm long, triangular. Ovary ovoid, ca. 1 mm, puberulent; style clavate, ca 1.5 mm long, conspicuously geniculate at base; stigma glabrous, with thickened lateral margins and a membranous apex. Capsules puberulent, green ovoid to oblongoid, ca. 4 – 7 mm long. Seeds dark brown, ovoid, ca. 1 mm long, glabrous.

#### Phenology

Chasmogamous flowers in April, cleistogamous flowers from May to September, and fruits from April to September.

#### Distribution and habitat

*Viola fluvialis* is currently only known from streamside habitats in Tongren Grand Canyon.

#### Conservation status

Due to the lack of further surveys in surrounding areas, the conservation status of this species is assessed as Data Deficient (DD) according to the IUCN criteria

***Viola orientosinensis*** Yan S. Huang, Y. Xiong & Q. Fan, sp. nov. (Fig. 10, 11)

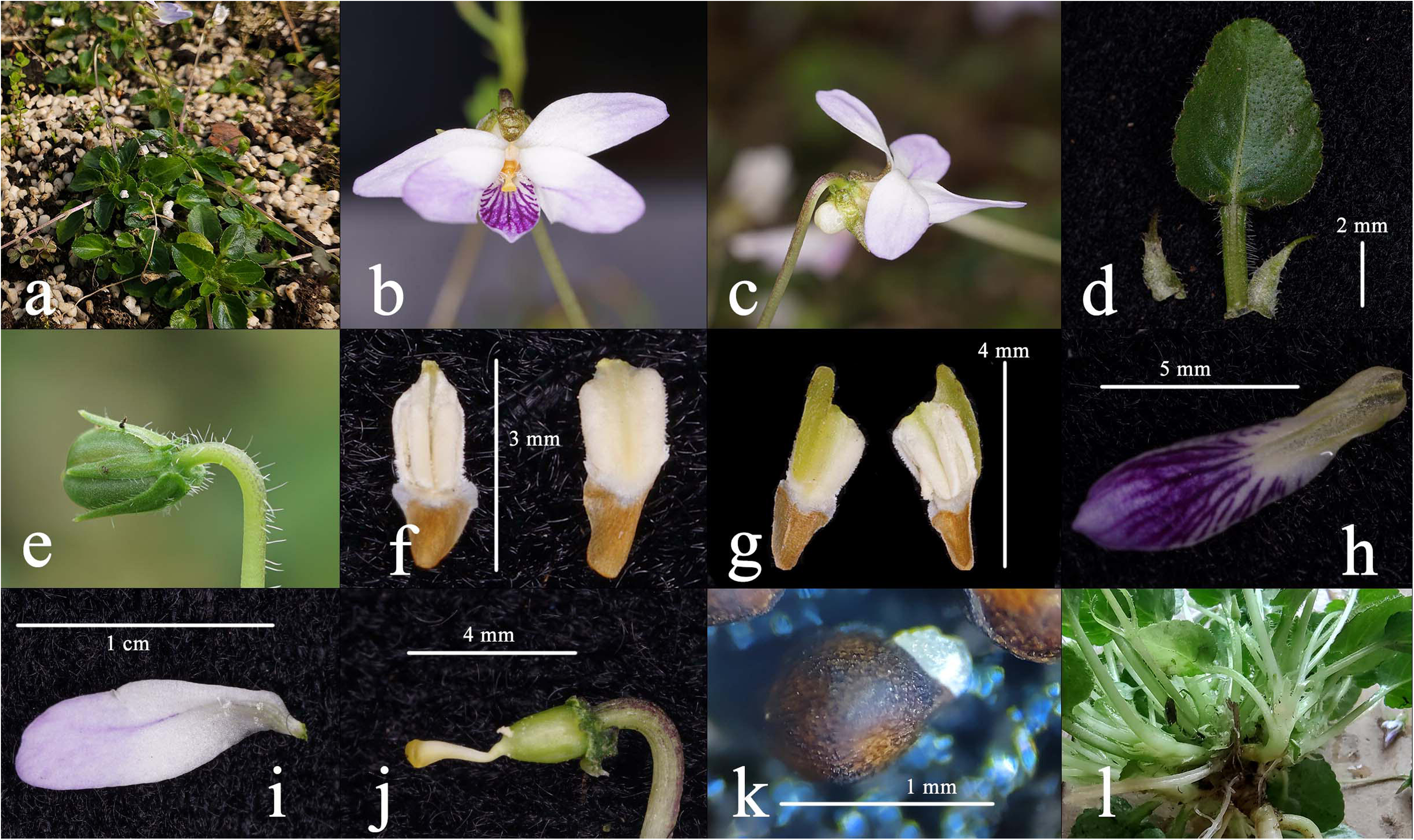

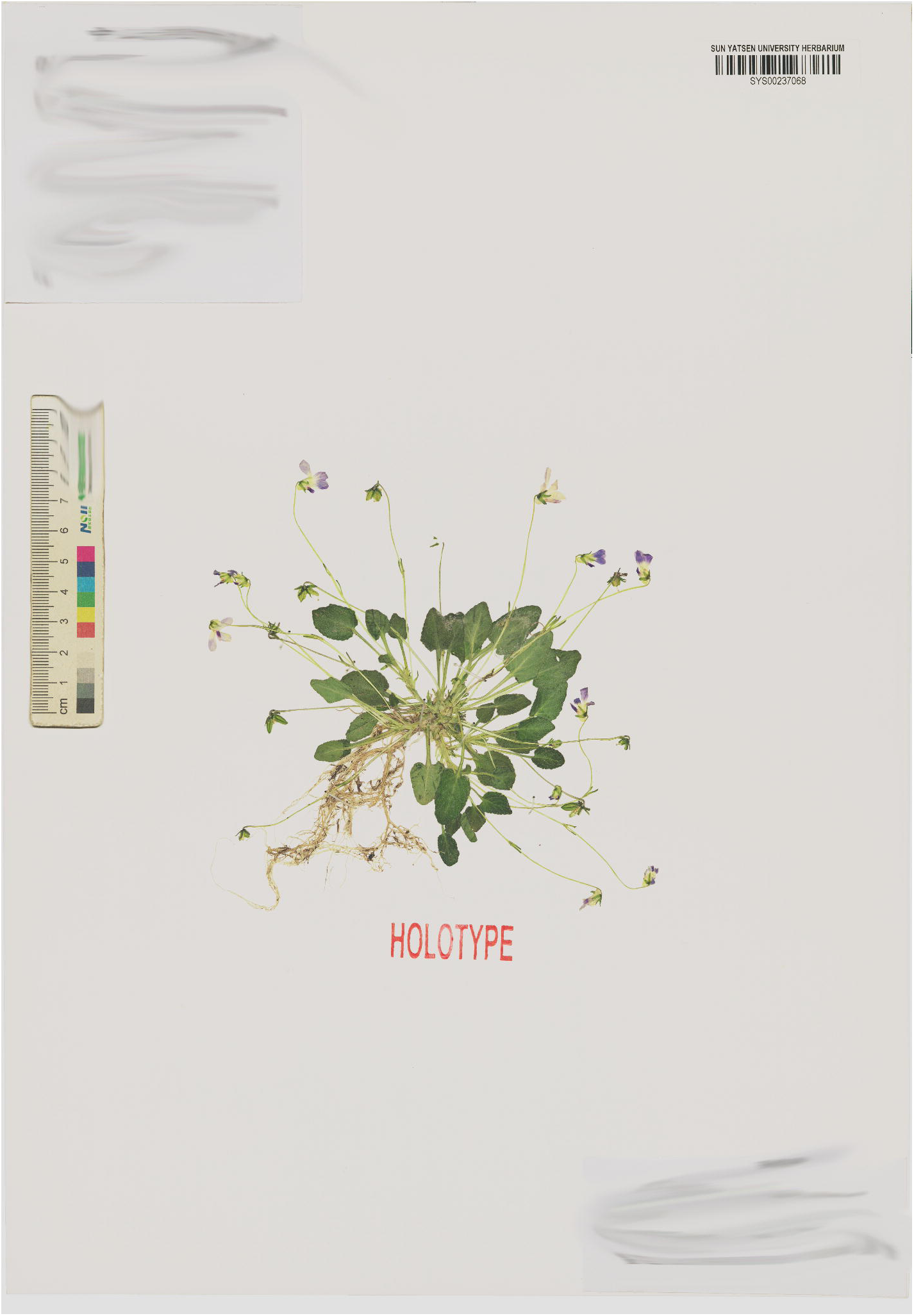

#### Diagnosis

A species morphologically close to *Viola nagasawae* Makino & Hayata but differing in having lateral petals bearded and lacking yellow patches at the base (vs. glabrous lateral petals with conspicuous yellow pacthes), leaf base distinctly decurrent to the petiole (vs. base not or only slightly decurrent), vegetative reproduction by tillering and lacking stolons(vs. by obvious stolons), robust taproot (vs. fibrous root system).

***Type*:** China. Jiangxi: Fuzhou City, Zixi County, Matoushan National Nature Reserve, in rocky crevices near the valley, 27°43′38.42″ N, 117°12′20.44″ E, 685 m a.s.l., 4 April 2024, *Yu Xiong Mts20240401* (holotype: SYS00237068; isotype: SYS00237065).

#### Etymology

The specific epithet refers to its distribution in eastern China

#### Description

Perennial, tillering herb with basal leaves rosulate, ca. 10 cm tall at flowering and ca. 3 cm tall when not flowering (lying on the ground). Rhizome short and robust, with dense remains of stipules and petioles. Stolons absent; lateral stems produced by tillering, each forming a rosette of leaves at the apex. Stipules 3 – 5 mm, adnate to petioles for about 1/3 at base, linear–triangular, apex acute, pubescent along the entire margin. Petioles 1.0 – 3.5 cm, with narrow wings, sparsely pubescent along the margin. Leaf blades ovate 1.1 – 1.3 × 1.4– 1.5 cm, sparsely pubescent along veins and margin; margin finely crenate; base cordate while living; apex obtuse. Chasmogamous flowers ca 1.2 cm in diam.; peduncles slender, 6 – 11 cm long, glabrous, rising well above the leaves, with two opposite bracteoles above middle; bracteoles linear-lanceolate, ca. 5 mm long, obtuse at apex, sparsely ciliate at base. Sepals green, with densely purple spots, puberulent, lanceolate, 0.8 – 1× 2 – 3 mm, with sparsely pubescent margin, apex obtuse, base truncate or rounded with extremely short semicircular appendages. Petals purple, the anterior one with apparent violet lines; posterior petals narrowly ovate, 3.5 × 8.5 mm, glabrous, with entire margin and obtuse apex; lateral petals with straight to slightly clavate hairs at the base, oblong, 4.0 × 9.0 mm, with entire margin and obtuse or erose apex; anterior petal oblong, with a short saccate spur at base, with interior side of base puberulent, including spur 0.9 cm long, with entire margin and acuminate apex. Stamens 5, unequal, puberulent; anther thecae 1.5 – 1.8 mm long, with terminal appendages ca 1.0 mm long; posterior appendages (nectar spurs) of the two anterior stamens ca 2.0 mm long, triangular. Ovary ovoid to ellipsoid, ca. 1.8 mm, puberulent; style clavate, ca 2.0 mm long, conspicuously geniculate at base; stigma glabrous, with thickened lateral margins and a membranous apex. Cleistogamous flowers in June ca. 5 mm long; peduncles 4 – 6 cm long; bracteoles linear-lanceolate, ca. 4 mm long, acuminate at apex. Sepals green, lanceolate, 0.9–1.2 × 2.8 – 4.0 mm, with entire margin and truncate base, acuminate at apex. Without conspicuous petals. Stamens 3, equal, 1.2 – 2.1 mm; terminal appendages triangular, ca. 0.8 mm. Capsules puberulent, light green to brown, some with purple spots at maturity, ovoid to oblongoid, ca. 7 mm long. Seeds brown, nearly globose, ca. 1 mm long, with small tubercles and conspicuous elaiosomes.

#### Phenology

Chasmogamous flowers from March to April, cleistogamous flowers from May to September, and fruits from April to November.

#### Distribution and habitat

*Viola orientosinensis* is currently known from Matoushan Nature Reserve in the Wuyi Mountains, as well as from Huangshan City, Anhui Province, and Hangzhou City, Zhejiang Province, eastern China. The species grows on rocks at altitudes of 600–1800m a.s.l. and can occur above the snowline on Huangshan Mountain.

#### Conservation status

Given its broad distribution, the species is evaluated as Least Concern (LC) according to the IUCN criteria(B).

***Viola suborbiculata*** Yan S. Huang & Q. Fan, sp. nov. (Fig. 12, 13)

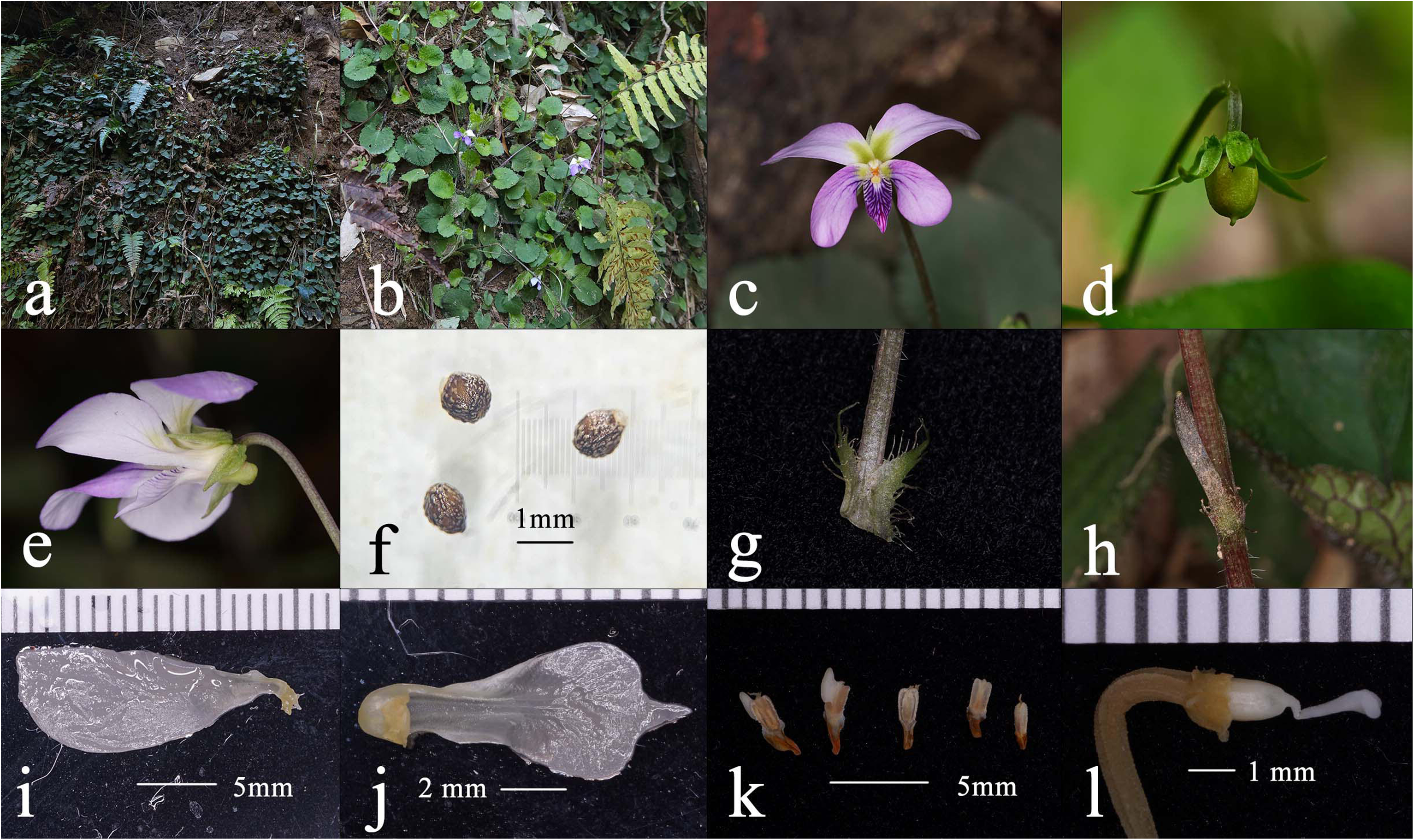

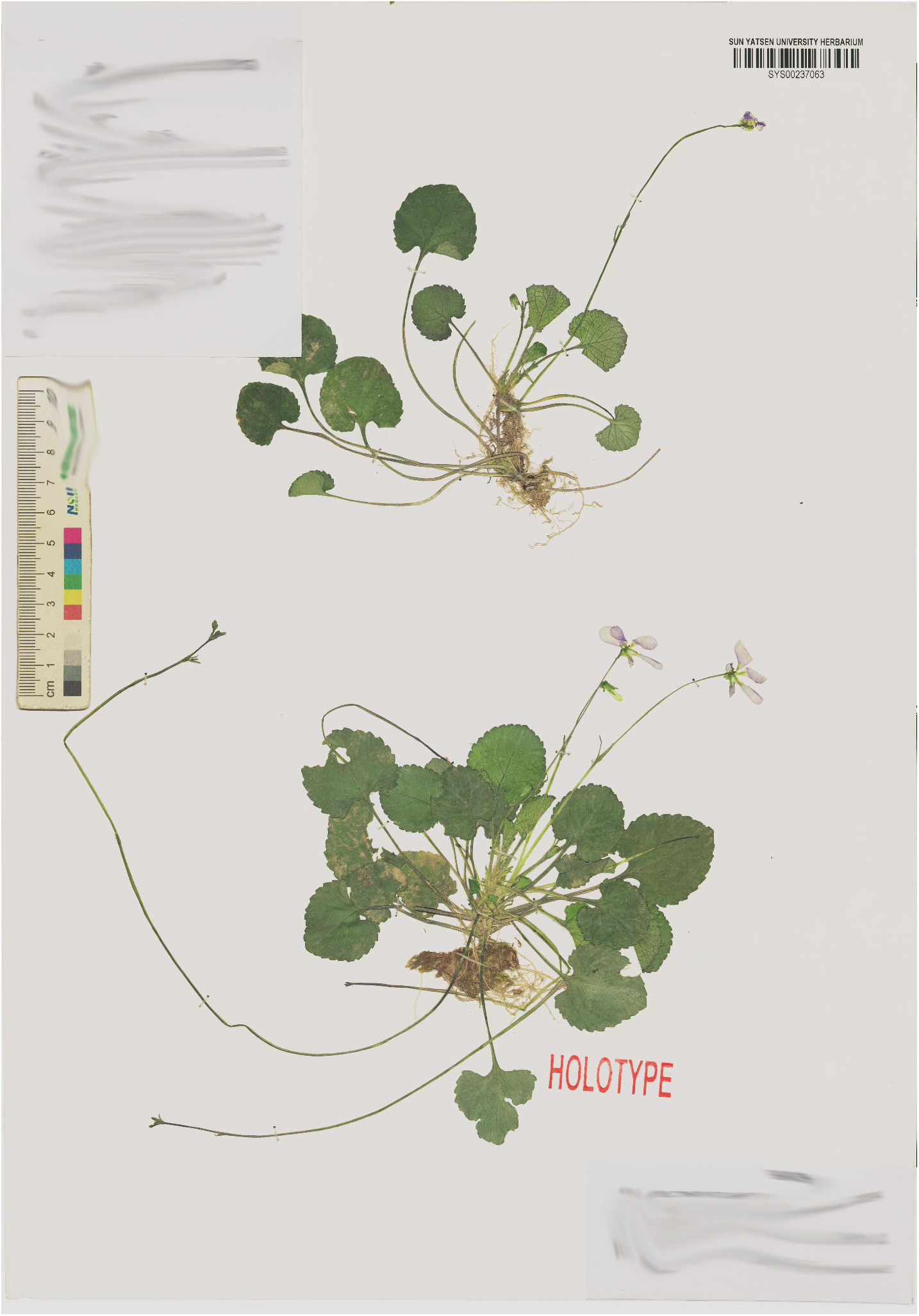

#### Diagnosis

This species is morphologically close to *Viola apoensis* Elmer. but differs in having larger leaves (2.5–3.0 × 2.0–3.0 cm vs. 1.0–1.5 × 0.75–1.0 cm), with serrate margins (vs. round-crenate). Flowers are larger, and the species has well-developed stolons (vs. stolons absent on the type specimens). The seeds are brown and with conspicuous tubercles (vs. lightly yellow and glabrous seeds).

***Type*:** China. Jiangxi: Yichun City, Yifeng County, Guanshan National Nature Reserve, on red-brown soils at forest edges, 28°33′22.4″ N, 114°33′40″ E, 368m a.s.l., 18 Mar 2025, *Yan. S. Huang HYS25031801*(holotype: SYS00237063; isotype: SYS00237064, SYS00237061, SYS00237062).

***Additional specimens examined:*** China. Jiangxi, Jiujiang, 23 May 2014, *Xuanhuai Zhan* et *Yansong Peng LPX1111* (LBG00138205); Hunan, Yueyang, 23 May 2015, *Xuanhuai Zhan et Yansong Peng LPX3190* (LBG00138211).

#### Etymology

The specific epithet refers its suborbicular leaf shape.

#### Description

Annual herb with basal leaves rosulate, ca. 10 cm tall at flowering and ca. 5 cm tall when not flowering. Rhizomes erect or obliquely erect, robust, with dense remains of stipules and petioles, 2.5 – 3.0 cm long. Lateral stolons well developed, spreading and elongating, usually producing adventitious roots with an apical rosette of leaves, and producing one-flowered pedicels at the nodes in the leaf axils. Stipules 3 – 5 mm, adnate to petioles for about 1/3 – 1/2 at base, triangular, acute at apex, with margins pinnatifid, glabrous. Petioles 5 – 8 cm, with dense purple spots, pubescent along the margin, narrowly winged near the blade base, the wings decurrent and gradually tapering toward the lower part. Leaf blade orbicular to suborbicular, 2.5 – 3.0 × 2.0 – 3.0 cm, dark green adaxially, pale green and nearly glabrous abaxially, sparsely pubescent along veins and margin; margin serrate; base cordate; apex obtuse. Chasmogamous flowers ca 2.5 cm in diam.; peduncles slender, 9 –14.5 cm long, pubescent on the lower half and glabrous above, rising well above the leaves, with two opposite bracteoles above middle; bracteoles linear-lanceolate, 0.7 – 1.0 cm long, acuminate at apex, densely ciliate at base. Sepals green, puberulent, lanceolate, 1.8 × 7 mm, with sparsely pubescent margin, apex acuminate, base truncate with short triangular appendages. Petals purple, the anterior one with apparent violet lines, the posterior and lateral petals with yellow to green spots at the base; posterior petals ovate, 1.1 × 1.7 cm, glabrous, with entire margin and obtuse apex; lateral petals with straight to slightly clavate hairs at the base, oblong, 0.8 × 1.9 cm, with entire margin and erose apex; anterior petal spathulate, expanded apically and acuminate, with a short saccate spur at base, base puberulent on the inner side, total length including spur 1.2 cm, margin entire. Stamens 5, unequal, puberulent; anther thecae 1.8 mm long, with terminal appendages ca 1.2 mm long; posterior appendages (nectar spurs) of the two anterior stamens 2.4 mm long, triangular. Ovary ovoid to ellipsoid, 2.5 mm long, glabrous; style clavate, 2.5 mm long, conspicuously geniculate at base; stigma glabrous, with thickened lateral margins. Cleistogamous flowers in June 4 –7 mm long; peduncles 2–5 cm long; bracteoles linear-lanceolate, ca. 4 mm long, apex acute. Sepals green, lanceolate, 1.4 –1.8 × 4.5 – 5.2 mm, with entire margin and truncate base, acuminate at apex. Without conspicuous petals. Stamens 2, equal, ca. 2 mm; terminal appendages triangular, ca. 0.8 mm. Capsules glabrous, green, ovoid to oblongoid, 4 – 10 mm long. Seeds brown, nearly globose, ca. 1 mm long, with obviously tubercles and very small elaiosomes.

#### Phenology

Chasmogamous flowers from March to May, cleistogamous flowers from May to September, and fruits from May to October.

#### Distribution and habitat

*Viola suborbiculata* is currently known from Guanshan National Nature Reserve in the Yichun City, Jiangxi Province as well as from Jiujiang City, Jiangxi Province, and Yueyang City, Hunan Province, China. The species grows at altitudes of 350 – 900m a.s.l.

#### Conservation status

Given its broad distribution, the species is evaluated as Least Concern (LC) according to the IUCN criteria(B).

***Viola tenuifolia*** Yan S. Huang et Q. Fan, sp. nov. (Fig. 14, 15)

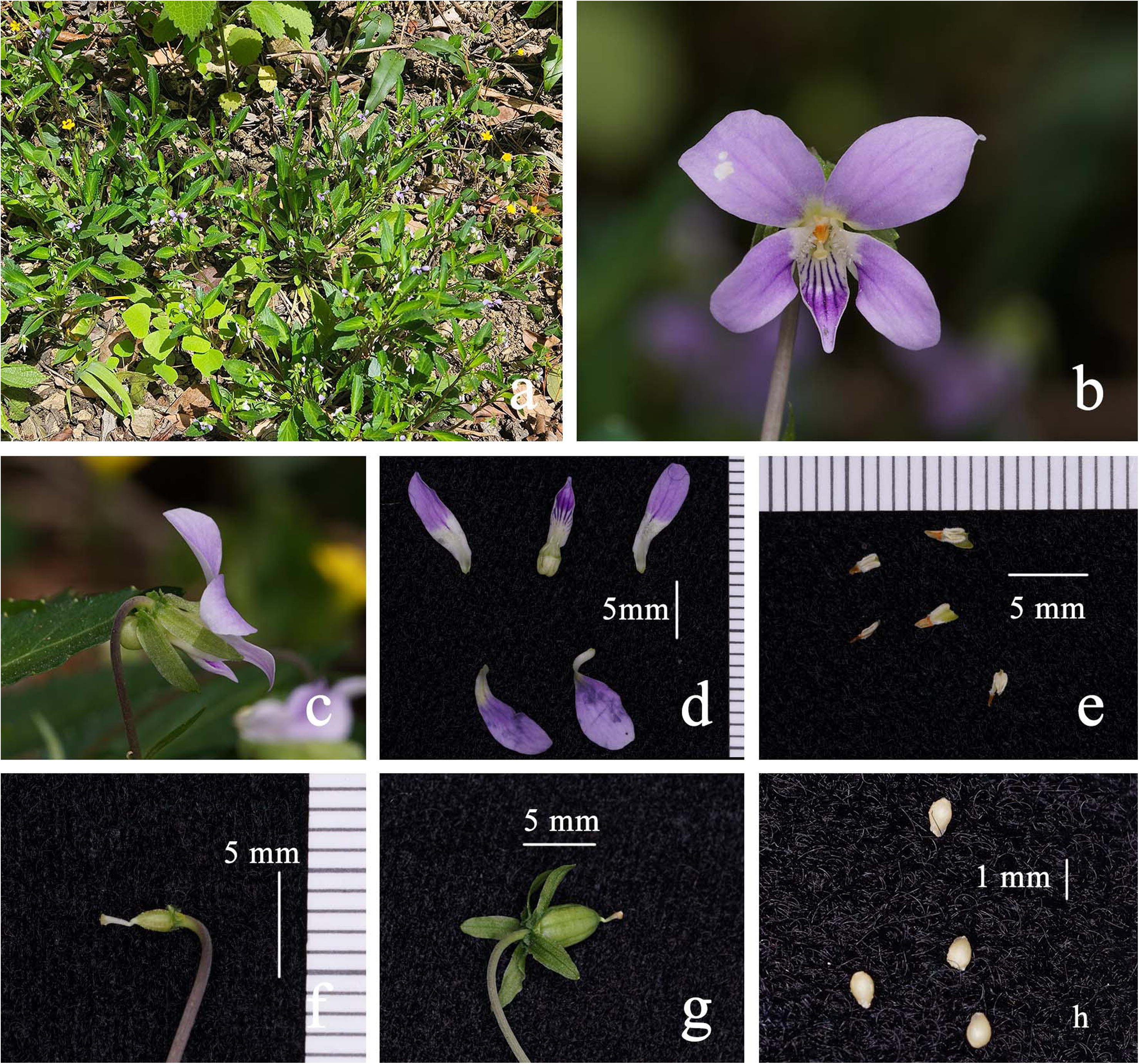

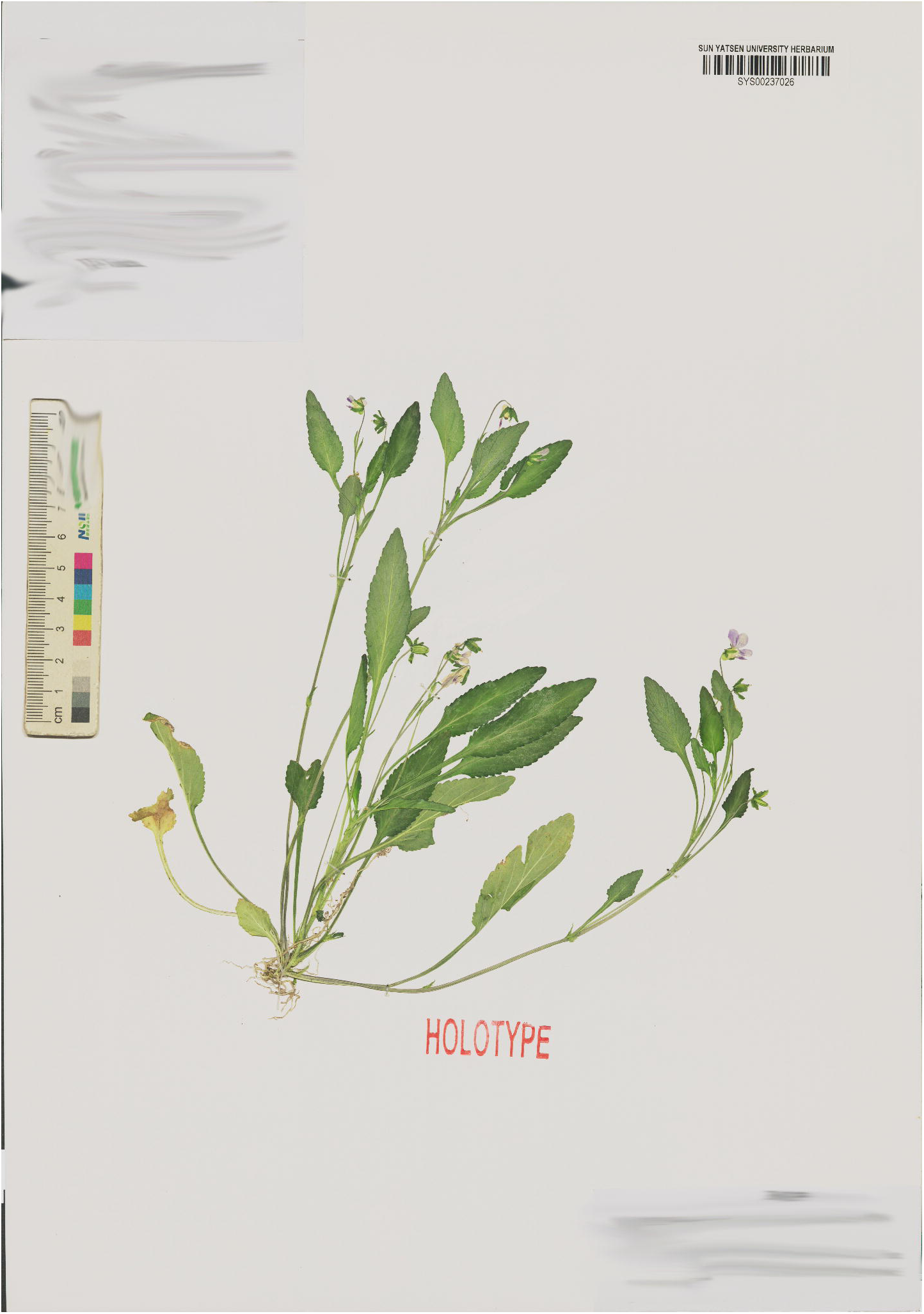

#### Diagnosis

A species molecularly close to *Viola diffusa*, but differing in having aerial stems and lateral stems that are erect or ascending (vs. lacking erect stems, with creeping lateral stolons); leaves lanceolate and nearly glabrous (vs. ovate and pubescent); seeds with tubercles (vs. smooth).

***Type:*** China. Hubei: Yichang City, Yiling District, in rocky crevices near the valley, 30°47′28.91″ N, 111°15′24.32″ E, 190 m a.s.l., 14 April 2025, *Yan. S. Huang HYS25041402* (holotype: SY00237026; isotype: SYS00237028).

***Additional specimens examined:*** China. Sichuan, Mount. Emei, 6 April 2014, *Lei Wu 4472* (BNU0039701); Chongqing, Pengshui, 24 July 1988, *Fading Fu* et *Yaling Cao 0219*(PE01356175); Chongqing, Pengshui, 24 July1988, *Fading Fu* et *Yaling Cao 0219*(CDBI0080716); Sichuan, Hongya, *Zewei Wang 648* (CDBI0080738); Sichuan, Hongya, *Zewei Wang 648* (CDBI0080740); Sichuan, Hongya, 7 July 1993, *Weikai Bao1629* (CDBI0080739); Sichuan, Hongya, *Zewei Wang 648* (CDBI0080740); Sichuan, Hongya, 7 July 1993, *Weikai Bao1629* (CDBI0080741).

#### Etymology

The specific epithet refers to its lanceolate leaves.

#### Description

Perennial, basal leaves rosulate, 18 – 20 cm tall with erect aerial stems. Rhizome short, ca. 2 cm. Stolons absent; lateral stems erect or ascending, ca. 20 cm tall, leaves produced at the nodes, with flower buds forming in the leaf axils. Stipules 6 – 7 mm, adnate to petioles at base, linear–lanceolate, apex acute, margin fimbriate, glabrous. Petioles 2.5–5 cm long, the leaf base decurrent along the petiole forming narrow wings, which are sparsely serrate in the upper part; glabrous. Leaf blades lanceolate, 1.2 – 1.7 × 4.6 – 5.2 cm, apex acuminate, margin serrate, base truncate or decurrent. Chasmogamous flowers 0.8 – 1.0 cm in diam.; Peduncles 5 – 11 cm long, glabrous; with a pair of bracteoles at about 1/2 or on the upper part; bracteoles linear-lanceolate, 4 – 7 mm long, apex acute, glabrous. Sepals green, glabrous, lanceolate, 1.5 × 6 mm, apex acuminate, base with short claw-like appendages. Petals purple, the anterior one with apparent violet lines; posterior petals obovate, 4 × 8 mm, glabrous, with entire margin and obtuse apex; lateral petals with straight to slightly clavate hairs at the base, oblong, 3 × 10 mm, with entire margin and obtuse or erose apex; anterior petal narrowly oblong, with a short saccate spur at base, and interior side of base puberulent, including spur 9 mm long, with entire margin and acuminate apex. Stamens 5, unequal, puberulent; anther thecae 1.5 mm long, with terminal appendages 0.5 – 1.0 mm long; posterior appendages (nectar spurs) of the two anterior stamens 1.2 mm long, triangular. Ovary ovoid, ca. 2 mm, puberulent; style clavate, ca 2.0 mm long, conspicuously geniculate at base; stigma glabrous, with thickened lateral margins and a membranous apex. Capsules puberulent, green ovoid to oblongoid, 4 mm long. Seeds brown, ovoid, ca. 1 mm long, with small tubercles, elaiosomes absent.

#### Phenology

Chasmogamous flowers in March, cleistogamous flowers from May to September, and fruits from April to September.

#### Distribution and habitat

*Viola tenuifolia* is currently known from Yiling Gorge in Hubei Province and grows in stream valleys. Otherwise, this species is widespread in Sichuan Province and Chongqing.

#### Conservation status

Given its broad distribution and having many populations, the species is evaluated as Least Concern (LC) according to the IUCN criteria(B)

***Viola wilsonii*** W.Becker, Bull. Misc. Inform. Kew 1928(6): 251. stat. rev. (Fig. 16)

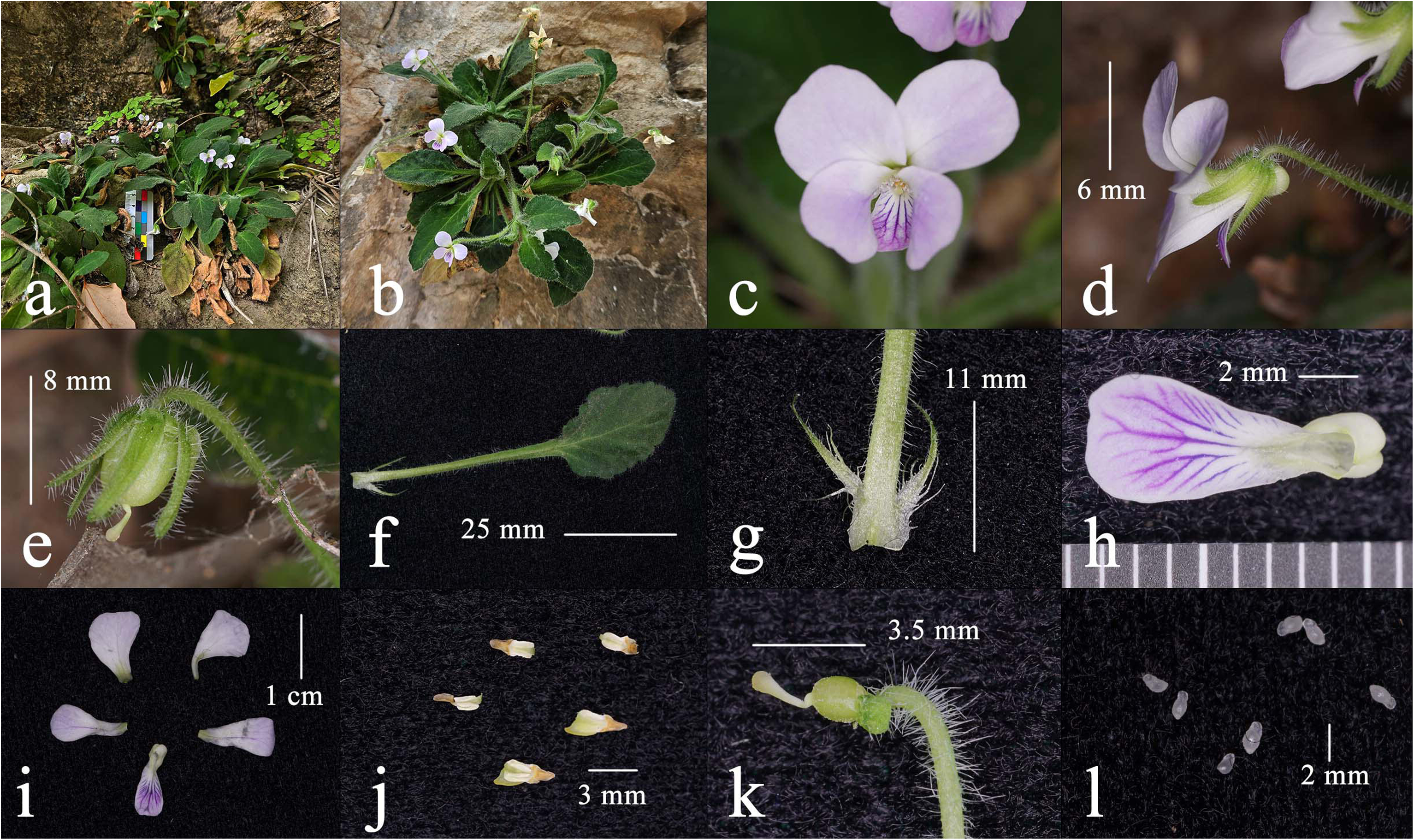

≡ *V. diffusa* var. *tomentosa* W. Becker, Beih. Bot. Centralbl. 20(2): 127. 1906.

#### Diagnose

This species is molecularly closely related to *Viola diffusoides*, but differs by the anterior petal apex being shallowly emarginate, the entire plant densely tomentose, and having a robust rhizome.

***Type*:** *China*: Western Hubei, 1900, *E.H. Wilson* 245 (K000254178).

***Additional specimens examined:*** China: Hubei, Yichang, 14 April 2025, *Yan. S. Huang HYS25041401* (SYS00237032).

#### Etymology

The specific epithet in memory of E.H. Wilson, the eminent plant collector who first gathered material of this species in western Hubei, China, in 1900.

#### Description

Perennial herb, ca. 8 cm tall, tomentose, with basal leaves tufted. Rhizomes erect or obliquely erect with very short internodes, robust, with dense remains of stipules and petioles, 2.5 – 3.0 × 0.4 –1 cm. Lateral stolons creeping or obliquely ascending, usually producing adventitious roots with an apical rosette of leaves. Stipules 10 – 12 mm, adnate to petioles for about 1/3 at base, triangular, acute at apex, with margins pinnatifid, pubescent. Petioles 1.5 – 5 cm, densely pubescent, with leaf base gradually decurrent into winged petiole. Leaf blade ovate to oblong, 1.4 – 3.8 × 0.8 – 1.9 cm, tomentose, margin serrate, apex acuminated. Chasmogamous flowers ca 1.5 cm in diam; peduncles 4.5 – 5 cm long, densely pubescent, with two opposite bracteoles above middle; bracteoles linear-lanceolate, 0.7 cm long, acuminate at apex, densely pubescent. Sepals green, densely pubescent, lanceolate, 5.0 – 6.5 mm long, apex acuminate, base truncate with extremely short orbicular appendages. Petals purple, the anterior one with apparent violet lines, the posterior and lateral petals with yellow to green spots at the base; Posterior petal obovate, 0.6 × 0.7 cm, glabrous, margin entire; apex very shallowly emarginate; lateral petals with straight to slightly clavate hairs at the base, oblong, 0.5 × 0.9 cm, with entire margin and very shallowly emarginate apex; anterior petal spathulate, expanded apically and shallowly emarginate at apex, with a short saccate spur at base, base puberulent on the inner side, total length including spur 8.5 cm, margin entire. Stamens 5, unequal, puberulent; anther thecae 1.1 – 1.5 mm long, with terminal appendages 1 – 1.2 mm long; posterior appendages (nectar spurs) of the two anterior stamens 1.7 mm long, triangular. Ovary ovoid, 1.5 mm long, puberulent; style clavate, 2.0mm long, conspicuously geniculate at base; stigma glabrous, with thickened lateral margins. Capsules glabrous, green, ovoid, ca. 6 mm long. Seeds yellow, ovoid, ca. 1 mm long, glabrous, with very obvious elaiosomes.

#### Phenology

Chasmogamous flowers from April to May, and fruits from May to August.

#### Distribution and habitat

*Viola wilsonii* is currently known only from Xiling Gorge, Yichang City, Hubei Province, China, where it grows on cliffs at an elevation of ca. 200 m.

#### Conservation status

*Viola wilsonii* is currently known from a single locality, despite intensive field surveys. Fewer than 50 individuals have been observed on cliffs in Xiling Gorge. According to the IUCN Red List Categories and Criteria (IUCN 2025), this species should be assessed as Critically Endangered (CR) under criterion D, due to its extremely small population size and highly restricted distribution.

#### Notes

*Viola wilsonii* W.Becker is here resurrected from synonymy under *V. diffusa* Ging. It was originally published by Becker as *V. diffusa* var. *tomentosa* W.Becker (1906), but later elevated by himself to species rank as *V. wilsonii*(1928).

According to the principle of priority and typification, the correct name for this taxon would have been *V. diffusa* var. *tomentosa* if it indeed belonged within *V. diffusa*. However, both morphological and molecular evidence indicate that this taxon is not a variety of *V. diffusa*, but more closely related to *V. diffusoides*. Therefore, it is here treated as a distinct species. The epithet “*tomentosa*” cannot be used at species rank, as it is already occupied by *V. tomentosa* M.S.Baker & J.C.Clausen (Leafl. W. Bot. 5: 142. 1949). Hence, the valid name for this species remains *V. wilsonii* W.Becker.

## Discussion

Molecular analyses based on ITS and GPI gene sequences provided clear insights into the phylogeny within *Viola* subsect. *Diffusae*. The ITS data primarily clarified the broader phylogenetic placement of the species within the genus and its major subsections, while the GPI data offered additional resolution for relationships at the subsection and species levels. ITS tree and the CHAM subgenome clade of the GPI gene supported several key clades, including the grouping of *V. diffusoides*, *V. wilsonii*, *V. aromatica*, and *V. tenuifolia*, as well as the monophyly of the *V. lucens*–*V. changii*–*V. acidophila* group. The GPI analyses revealed additional details on species relationships, such as the close affinity of *V. orientosinensis* with *V. qingruii* and *V. amamiana*, rather than with *V. nanlingensis* as indicated in the ITS tree. *V. nanlingensis* was found to be non-monophyletic, although a clear phylogenetic affinity was observed between this species and *V. suborbiculata*. *V. acidophila* clustered together with specimens of *V. changii* from different localities, with *V. changii* forming a monophyletic group; these two species were recovered as well-supported sister clades (bootstrap = 100; 100). And the systematic position of *V. fluvialis* showed substantial differences between the two GPI subgenomes.

The incongruent placement of species between the two GPI subgenomes, as well as between the GPI and ITS trees, suggests a complex evolutionary history potentially involving ancient hybridization or incomplete lineage sorting. Such conflicts between subgenomes have also been reported in other *Viola* lineages (Marcussen et al. 2022).

Morphological and molecular phylogenetic data together provide complementary evidence that informs a comprehensive view of species relationships in this geuns. In some cases, morphological differentiation corresponds well with molecular phylogenetic patterns. For example, *V. acidophila* is recovered as sister to a monophyletic *V. changii* in the GPI tree and is morphologically distinct in having oblong, densely puberulent leaves with an acuminate apex and dull surface (vs. ovate, glabrous or roughly pubescent, glossy leaves in *V. changii*), as well as by lacking stolons and reproducing via tillering.

Sometimes morphological similarity does not always reflect phylogenetic relationships. For example, though *V. aromatica* resembles *V. diffusa* in leaf form, this species is phylogenetically closer to *V. tenuifolia*, sharing traits like narrow leaves, but differs in pubescence, flower size, and scent. *V. fluvialis*, though resembling *V. yunnanensis* in morphology, has conflicting placements between the two GPI subgenomes, suggesting a complex evolutionary history possibly involving whole-genome duplication.

The type specimens of *Viola diffusa* are collected from Nepal, with the accession numbers K00674037, K000254254, and K000254255. Careful examination of these specimens revealed that K000254255 differs markedly from the others: its leaf apex is obtuse, the leaf base is deeply cordate, and the basal extension of the leaf is relatively narrow, corresponding to the newly described species *V. qingruii*. Notably, this specimen had previously been marked with a pencil to indicate its distinctiveness. In contrast, the remaining specimens exhibit densely pubescent leaves, an extremely broad leaf base extending to the petiole, and well-develope diffusa d stolons. Comparison with Chinese specimens labeled as *V. diffusa* from Yunnan and Tibet (Appendix Figure 1) shows similarity to the type, with a shared distinctive feature of smaller flowers, consistent with earlier observations (Zhou et al., 2008).

*Viola diffusoides* and *V. wilsonii* were previously treated as synonyms of *V. diffusa*, but previous studies have demonstrated that this treatment is inappropriate. Morphologically, *V. diffusoides* is characterized by larger flowers, glabrous stems and leaves, and a leaf base that is typically truncate rather than extending basally. Although an unpublished thesis reported that this species develops pubescence under cultivation, we did not observe this phenomenon during years of cultivation; instead, putative hybridization with *V. diffusa* in the Beibei population may explain the reported occurrence of hairs. *V. wilsonii* was first collected in 1900. Its type specimen (K000254178) exhibits shallowly bifid petals, dense white puberulence throughout the plant, and thick, often highly developed rhizomes, with the original locality recorded only as western Hubei. Based on analysis of Ernest Henry Wilson’s diaries and field tracking, we ultimately located populations matching all morphological characteristics on cliffs in Yichang (Appendix Figure 2). Notably, the seeds of this species contain oil bodies, a feature similar to that of *V. pendulipes*, which also grows on karst landscapes (Huang et al., 2023b).

*Viola orientosinesis* is noted in our records as widely distributed in Jiangxi, Anhui, and Zhejiang, but no additional examined specimens are provided. Extensive hybridization occurs among *V. orientosinesis* and the two species *V. qingruii* and *V. nanlingensis*, resulting in *V. orientosinesis* exhibiting some intermediate traits, such as yellow basal spots on the petals, dense puberulence, and widened leaf bases (Appendix Figure 3). This hybridization can be confirmed by trait segregation in the offspring as well as by double peaks observed in Sanger sequencing. The population from Matoushan, Jiangxi, is the only one in which pure individuals can be reliably identified by Sanger sequencing, and their characteristics correspond to those described above. Additionally, we noted that *V. orientosinesis* and *V. acidophila* lack stolons, hence some may argue that this does not conform to the characteristics of the subsect. *Diffusae*; however, the earlier described *V. guangzhouensis* in the same subsect. often produces lateral stems as aerial stems rather than stolons. Thus, stoloniferous growth is a common but not absolute trait, and subsection classification should be based on monophyly, as discussed Marcussen et al., 2022.

### Key to subsect. *Diffusae*

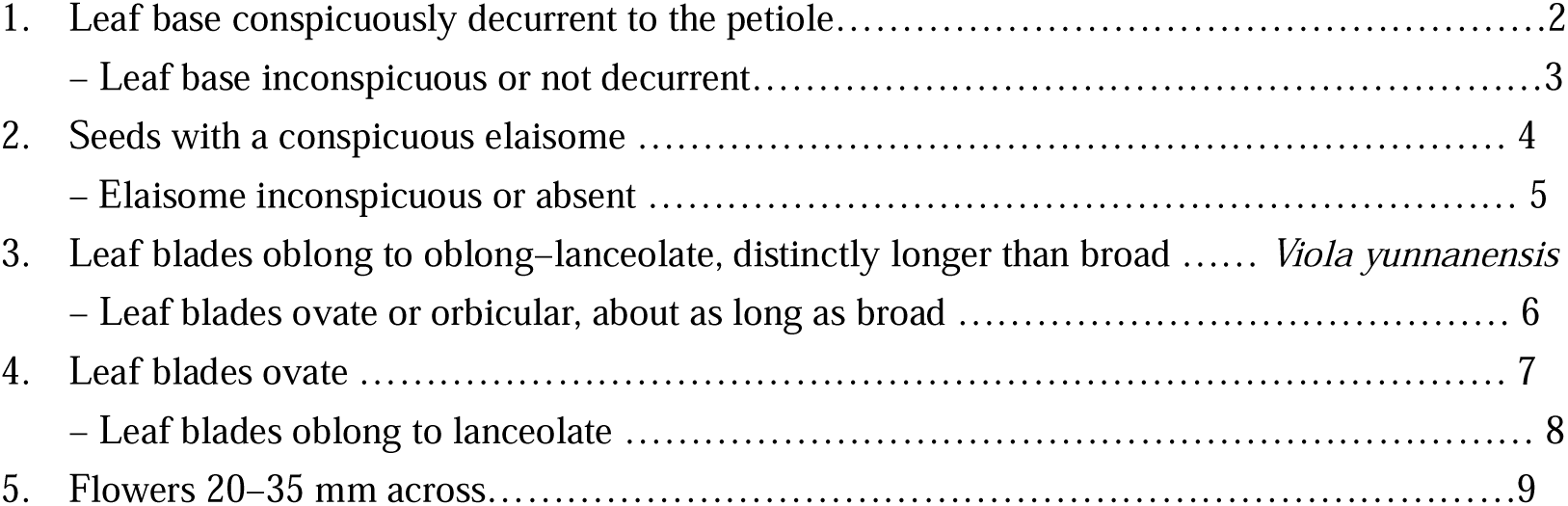
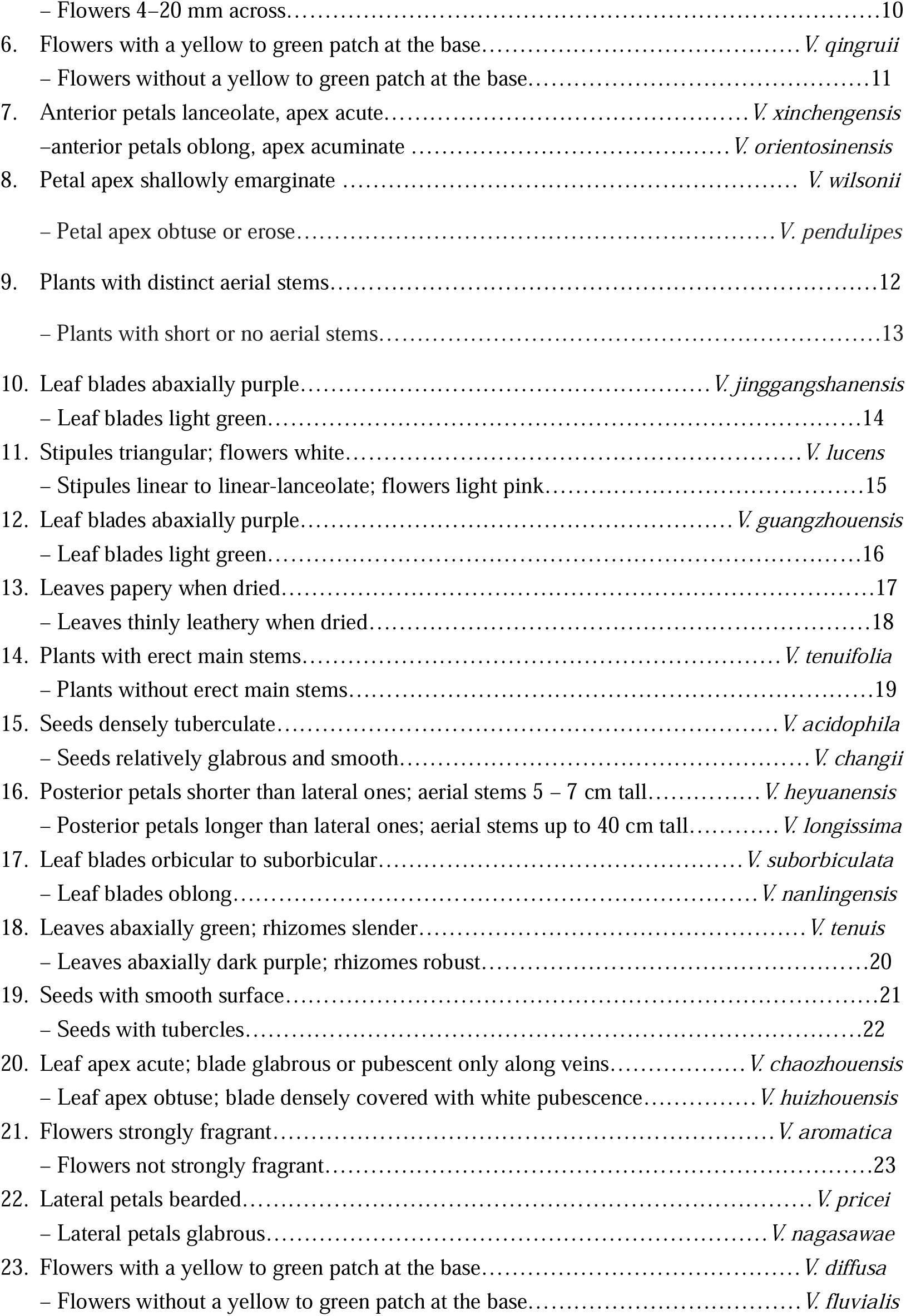

## Reference

Chen S., Zhou Y., Chen Y., Gu J. 2018. fastp: an ultra-fast all-in-one FASTQ preprocessor. –Bioinformatics 34: i884–i890.

Clausen, J. 1929. Chromosome number and relationship of some North American species of *Viola*. – Ann. Bot. 43: 741–764.

Danecek P., Bonfield J.K., Liddle J., et al. 2021. Twelve years of SAMtools and BCFtools. –Gigascience 10: giab008. 10.1093/gigascience/giab008

Doyle, J. J. and Doyle, J. L. 1987. A rapid DNA isolation procedure for small quantities of fresh leaf tissue. – Phytochem. Bull. 19: 11–15.

Fan, Q., Chen, S. F., Wang, L. Y., Chen, Z. X. and Liao, W. B. 2015. A new species and new section of *Viola* (Violaceae) from Guangdong, China. – Phytotaxa 197: 15–26.

Huang Y.S., Ding J.H., Ye Q.L., Dai J.M., Zhong Z.M., Fan Q. 2023a. Four new species of *Viola* (Violaceae) from southern China. – Nord. J. Bot. 41(6): e03941.

Huang, Y. S., Jia, X. Y., Zeng, Q. G., Wen, W. L. and Fan, Q. 2023b, *Viola pendulipes* (Violaceae), a new species from Guangdong Province, China. – Nord. J. Bot. 2023: e04165.

Katoh K., Standley D.M. 2013. MAFFT multiple sequence alignment software version 7: improvements in performance and usability. – Mol. Biol. Evol. 30: 772–780. 10.1093/molbev/mst010

IUCN. 2025. The IUCN Red List of Threatened Species. Version 2025–1. https://www.iucnredlist.org. Accessed on [24-07-2025].

Li H., Durbin R. 2009. Fast and accurate short read alignment with Burrows –Wheeler transform. – Bioinformatics 25: 1754–1760. 10.1093/bioinformatics/btp324

Li, X. C., Wang, Z. W., Wang, Q., Ge, B. J., Chen, B., Yu, P., Zhong, X. 2022. *Viola shiweii*, a new species of *Viola* (Violaceae) from karst forest in Guizhou, China. – PhytoKeys. 196:63–89. 10.3897/phytokeys.196.83176

Marcussen, T., Ballard, H. E., Danihelka, J., Flores, A. R., Nicola, M. V. and Watson, J. M. 2022. A revised phylogenetic classification for Viola (Violaceae). – Plants 11: 2224.

Minh, B. Q., Schmidt, H. A., Chernomor, O., Schrempf, D., Woodhams, M. D., Von Haeseler, A. and Lanfear, R. 2020. IQ-TREE 2: new models and efficient methods for phylogenetic inference in the genomic era. – Mol. Biol. Evol. 37: 1530 –1534.

Ning, Z. L., Zeng, Z. X., Chen, L., Xu, B. Q. and Liao, J. P. 2012. *Viola jinggangshanensis* (Violaceae), a new species from Jiangxi, China. – Ann. Bot. Fenn. 49: 383–386.

Tamura, K., Stecher, G., Peterson, D., Filipski, A., and Kumar, S. 2013. MEGA6: molecular evolutionary genetics analysis version 6.0. – Mol. Biol. Evol. 30: 2725–2729.

White, T. J., Bruns, T., Lee, S. J. W. T. and Taylor, J. 1990. Amplification and direct sequencing of fungal ribosomal RNA genes for phylogenetics. – In: Innis, M. A., Gelfand, D. H., Sminsky, J. J. and White, T. J. (eds), PCR protocols: a guide to methods and applications, vol. 18. pp. 315–322.

Zhou, J.S., Gong, Q., Xing, F.W. 2008. *Viola nanlingensis* (Violaceae), a new species from Guangdong, southern China. – Ann. Bot. Fenn. 45(3): 233–236.

