## Supplemental table 1 for "Six New Species and Two Reinstatements of *Viola* (Violaceae) from China"

Table S1. GenBank accessions for phylogenetic analysis

| *V. biflora* | DQ055348 |
| --- | --- |
| *V. canadensis* | AF097231,  MG234951 |
| *V. pubescens* | DQ006044 |
| *V. sempervirens* | MG235908 |
| *V. sheltonii* | AF097226,  AF097272 |
| *V. uniflora* | AY582167,  AY541600 |
| *V. urophylla* | MH117805 |
| *V. austrosinensis* | OM406228 |
| *V. kwangtungensis* | OM406230 |
| *V. mucronulifera* | FJ002910 |
| *V. sumatrana* | OM406231 |
| *V. raddeana* | AY928279 |
| *V. triangulifolia* | FJ002912 |
| *V. amamiana* | JF830899 |
| *V. diffusa* | MH711723 |
| *V. guangzhouensis* | MW683480 |
| *V. huizhouensis* | MW683486 |
| *V. lucens* | FJ002913 |
| *V. nanlingensis* | FJ002916 |
| *V. yunnanensis* | FJ002915 |
| *V. chaerophylloides* | DQ787762 |
| *V. albida* | AY928292 |
| *V. dissecta* | JQ950564 |
| *V. patrinii* | AY928298 |
| *V. selkirkii* | AY928307 |
| *V. somchetica* | HM851457 |
| *V. tashiroi* | JF830885 |
| *V. variegata* | KC330743 |
| *V. epipsila* | MG237736 |
| *V. grandisepala* | FJ002903 |
| *V. lanceolata* | MG235616 |
| *V. minuscula* | AF097236,  AF097282 |
| *V. moupinensis* | FJ002900 |
| *V. palustris* | KX166144 |
| *V. principis* | FJ002904 |
| *V. yazawana* | AY928289 |
| *V. heyuanensis* | OP935140 |
| *V. chaozhouensis* | OP935142 |
| *V. qingruii* | OP935150 |
| *V. tenuis* | OP935156 |
| *V. longissima* | OP935160 |
| *V. yunnanensis* | FJ002915 |
| *V. eizanensis* | DQ787773 |
| *V. woosanensis* | KP227493 |
| *V. violacea* var. *violacea* | LC527435 |
| *V. sieboldii* | LC643088 |
| *V. tokubuchiana* var. *takedana* | AB759264 |
| *V. iwagawai* | JF830880 |
| *V. dactyloides* | JQ950563 |
| *V. phalacrocarpa* | AY928294 |
| *V. pinnata* | JQ950572 |
| *V. dactyloides* | JQ950563 |
| *V. yunnanfuensis* |  |
| *V. changii* |  |
| *V. pilicalcarata* |  |
| *V. orientosinensis* |  |
| *V. acidophila* |  |
| *V. fluvalis* |  |
| *V. suborbiculata* |  |
| *V. diffusoides* |  |
| *V. wilsonii* |  |
| *V. aromatica* |  |
| *V. tenuifolia* |  |
| *V. pendulipes* | OR483797 |
| *V. xinchengensis* | PV089292 |
